# Recurrent cancer-associated ERBB4 mutations are transforming and confer resistance to targeted therapies

**DOI:** 10.1101/2024.12.19.626686

**Authors:** Veera K. Ojala, Sini Ahonen, Aura Tuohisto-Kokko, Olaya Esparta, Peppi Suominen, Anne Jokilammi, Iman Farahani, Deepankar Chakroborty, Nikol Dibus, Steffen Boettcher, Tomi T. Airenne, Mark S. Johnson, Lisa D. Eli, Klaus Elenius, Kari J. Kurppa

## Abstract

Receptor tyrosine kinase ERBB4 (HER4) is frequently mutated in human cancer, and *ERBB4* mutations have been identified in patients relapsing on targeted therapy. Here, we addressed the functional consequences of recurrent cancer-associated *ERBB4* mutations that are located at regions important for dimer interactions and/or are paralogous to known oncogenic hotspot mutations in other *ERBB* genes. Eleven out of 18 analyzed mutations were transforming in cell models, thus suggesting oncogenic potential for more than half of the recurrent ERBB4 mutations. More detailed analyses of the most potent mutations, S303F, E452K and L798R, showed that they are activating, can co-operate with other ERBB receptors and are targetable with clinically available second-generation pan-ERBB inhibitors neratinib, afatinib and dacomitinib. Furthermore, the S303F mutation, together with a previously identified activating ERBB4 mutation, E715K, promoted resistance to third-generation EGFR inhibitor osimertinib in EGFR-mutant lung cancer model *in vitro* and *in vivo*. Together, these results are expected to facilitate clinical interpretation of the most recurrent cancer-associated *ERBB4* mutations. The findings provide rationale for testing the efficacy of clinically used pan-ERBB inhibitors in patients harboring driver *ERBB4* mutations both in the treatment-naïve setting, and upon development of resistance to targeted agents.

## INTRODUCTION

All four members of the epidermal growth factor receptor (EGFR)/ERBB/HER family of receptor tyrosine kinases (RTK) are frequently mutated or amplified in cancer (Yarden and Pines, 2012; Arteaga and Engelman, 2014). Numerous ERBB-targeted therapies have been developed but their currently approved indications include only cancers with *EGFR* or *ERBB2* alterations. Of these, the second-generation irreversible ERBB tyrosine kinase inhibitors (TKI) neratinib, afatinib and dacomitinib, are termed pan-ERBB inhibitors as they potently inhibit EGFR, ERBB2 and ERBB4, although not directly the kinase-impaired ERBB3 (Davis *et al*., 2011). Yet, despite the high frequency of *ERBB4* missense mutations in various cancer types and characterization of several potentially oncogenic *ERBB4* mutations (Prickett et al. 2009; Nakamura et al. 2016; Chakroborty et al. 2022; Kurppa et al. 2016; Tvorogov et al. 2009), the rationale for clinically targeting ERBB4 in cancer has not been fully developed. The reasons for this include the perplexing results of the role of ERBB4 in cancer as well as the rarity of the distinct *ERBB4* mutations that have been characterized as oncogenic thus far.

*ERBB4* mutations have also been found in patients who developed resistance to EGFR-targeted therapies (Cremolini *et al*., 2019; Jänne *et al*., 2022; Vokes *et al*., 2022; Yaeger *et al*., 2023), warranting further investigation into the potential role of *ERBB4* mutations in therapy resistance. In addition, activation of ERBB4 signaling has been associated with EGFR- and ERBB2-targeted therapy resistance in multiple studies (Carrión-Salip *et al*., 2012; Canfield *et al*., 2015; Donoghue *et al*., 2018; Shi *et al*., 2018; Vokes *et al*., 2022; Debets *et al*., 2023), raising the possibility that mutant ERBB4 could be a potential therapeutic target to combat resistance to targeted therapies.

Unlike other ERBB receptors, ERBB4 has four different isoforms that have partly different, even opposing functions, and both tumor-suppressive and oncogenic functions have been described for ERBB4 (Määttä *et al*., 2006; Sundvall *et al*., 2010; Wali *et al*., 2014; Brockhoff, 2022; Lucas *et al*., 2022). Importantly, the oncogenic functions have been attributed to the ERBB4 isoforms that are predominant in cancer tissues and/or whose relative expression compared to the other isoforms is increased in cancer (Gilbertson *et al*., 2001; Junttila *et al*., 2005; Veikkolainen *et al*., 2011).

Another challenge in studying ERBB4 in cancer has been to choose which of the hundreds of different cancer-associated *ERBB4* mutations to select for functional analysis to identify potential oncogenic drivers, since *ERBB4* does not have clear mutational hotspots that would suggest functional relevance. Thus far, the strategies to characterize activating *ERBB4* mutations have included i) selecting mutations found in a specific cancer type, including melanoma and lung cancer (Prickett *et al*., 2009; Kurppa *et al*., 2016), ii) an unbiased functional screen of nearly all theoretically possible *ERBB4* missense mutations (Chakroborty *et al*., 2022), and iii) selecting *ERBB4* mutations found from cancer cell lines that are sensitive to ERBB-targeting drugs (Koivu *et al*., 2021). Although several activating gain-of-function *ERBB4* mutations have indeed been identified and many of these mutants have also been shown to be sensitive to pan-ERBB inhibitors, the rarity of these specific variants in patients makes it difficult to clinically assess their predictive value for ERBB4-targeted therapy.

The immensely increased body of tumor sequencing data over the past ten years has, however, resulted in the emergence of specific recurrent *ERBB4* mutations in hotspot-like regions of the *ERBB4* gene. Additionally, the rare but transforming *ERBB4* mutations structurally characterized thus far (Kurppa *et al*., 2016; Chakroborty *et al*., 2022) act via stabilizing dimerization interfaces similar to many of the known oncogenic hotspot mutations of ERBB2 and ERBB3 (Greulich *et al*., 2012; Jaiswal *et al*., 2013). To better facilitate the evaluation of the predictive potential of the cancer-associated *ERBB4* mutations, we functionally characterized 18 of the recurrent *ERBB4* variants. We show that majority (11/18) of the *ERBB4* mutations have significant transforming potential, and that the most potent mutants are biochemically activating and co-operate with other ERBB receptors - most strikingly S303F. We also demonstrate for the first time that the activating *ERBB4* mutations are able to promote resistance to EGFR-targeted therapy in the context of EGFR-mutant non-small cell lung cancer (NSCLC). Importantly, our data indicate that recurrent *ERBB4* gain-of-function mutations are targetable with clinically used second-generation pan-ERBB inhibitors.

## MATERIALS AND METHODS

### Cell culture

Phoenix-AMPHO (a gift from Dr. Garry Nolan), Ba/F3 (DSMZ) and PC-9 cells were cultured in RPMI 1640 (EuroClone, Gibco or Lonza), MCF10a cells (ATCC CRL-10317) in DMEM/F-12 (Lonza), and HEK-293T and COS-7 cells in DMEM (EuroClone, Gibco or Lonza). All media were supplemented with 10% FCS (Biowest), 50 U/ml penicillin and streptomycin solution (Gibco or Lonza) and 2 mM L-glutamine (Gibco or Lonza). Ba/F3 cell medium was additionally supplemented with 5% WEHI cell-conditioned medium as a source of IL3 while the media for IL3-independent Ba/F3 cells were supplemented or not with neuregulin-1β (NRG-1) (396-HB-050; R&D Systems). MCF10a cell medium was additionally supplemented with 20 ng/ml EGF (AF-100-15; Peprotech), 0.5 *µ*g/ml hydrocortisone (H-0888; Sigma), 100 ng/ml cholera toxin (C-8052; Sigma), and 10 *µ*g/ml insulin (I9278; Sigma). Cells were routinely tested for mycoplasma infection using MycoAlert (Lonza).

### Plasmids

The following previously described retroviral mammalian expression plasmids were used: *pBABE-puro-gateway*-*eGFP* (Chakroborty *et al*., 2022), *pBABE-puro-gateway-EGFR* (Chakroborty *et al*., 2019), *pBABE-puro-gateway-ERBB2* (Koivu *et al*., 2021), *pBABE-puro-gateway*-*ERBB3* (Koivu *et al*., 2021)*, pBABE-puro-gateway-ERBB4JM-aCYT-2* (Chakroborty *et al*., 2022)*, pMLDg/pRRE*, *pMD2.G*, and *pRSV-Rev* (gifts from Dr. Didier Trono; Addgene plasmids #12251, #12259, and #12253) (Dull *et al*., 1998). Expression constructs encoding ERBB4 JM-a CYT-2 point mutants listed in Supplementary Table S1 were created by cloning mutant inserts from *pDONR221*-vectors (Genscript) into *pBABE-puro-gateway* retroviral mammalian expression vector (a gift from Matthew Meyerson; Addgene plasmid #51070; http://n2t.net/addgene:51070; RRID:Addgene_51070)(Greulich *et al*., 2012) using LR clonase II mix (Invitrogen) for the LR Gateway recombination reaction. All generated constructs were verified by sequencing the insert.

### Generation of cell lines with stable ERBB4 expression

For retroviral transductions, Phoenix-AMPHO packaging cells were transfected with retroviral *pBABE-puro-gateway* constructs encoding ERBB4 variants or enhanced green fluorescent protein (eGFP) (vector control) using FuGENE 6 transfection reagent (Promega) according to manufacturer’s protocol. The retroviral supernatants of Phoenix-AMPHO cells were harvested 24 hours and 48 hours after transfection and used immediately to infect Ba/F3, MCF10a, or PC-9 cells. Cell pools with stable expression were selected with 2 *µ*g/ml puromycin (Gibco) for 48 hours and then maintained in 1 *µ*g/ml puromycin.

### Ba/F3 cell transformation assay

Ba/F3 cells stably expressing *ERBB4* variants or vector control were washed twice with PBS to deplete IL3 and 1×10^6^ cells were seeded in 10 ml IL3-free culture medium supplemented or not with 20 ng/ml NRG-1 or with IL3 (5% WEHI cell-conditioned medium) as a control. Cell viability was measured from day 0 up to saturation density with the MTT assay (CellTiter 96 nonradioactive cell proliferation assay; Promega) by collecting 100 *µ*l of the cell suspension into 96-well plate wells in quadruplicates. The doubling times were calculated using the equation 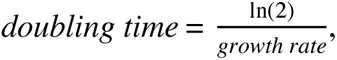 and 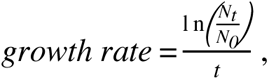 where, t is the time in hours reaching saturation density, *N_t_*is absorbance at saturation density, and *N_0_* is the absorbance at seeding density. The doubling times for cells growing in 20 ng/ml NRG-1 were normalized to the average of wild-type ERBB4 of each independent experiment. Kruskal Wallis test was used for statistical testing and false discovery rate was controlled by two-stage linear step-up method of Benjamini, Krieger and Yekutieli, considering *q* < 0.05 as significant. GraphPad Prism 10 was used to perform statistical analyses and creating the dot plots.

### MCF10a cell transformation assay

MCF10a cells stably expressing ERBB4 variants or vector control were seeded at 2,000 cells/well density into 96-well plate wells in triplicates in the presence of EGF. Next day, EGF-free medium supplemented or not with 50 ng/ml NRG-1 was changed and cell viability was measured with the MTT assay on days 0 and 8 of EGF deprivation. Day 8 cell viability fold changes to day 0 were calculated and normalized to the average of wild-type ERBB4 of each independent experiment. Kruskal Wallis test was used for statistical testing and false discovery rate was controlled by two-stage linear step-up method of Benjamini, Krieger and Yekutieli, considering *q* < 0.05 as significant. GraphPad Prism 10 was used to perform statistical analyses and creating the dot plots.

### Structural analysis

The S303F mutation was analyzed using the crystal structure PDB:2AHX (2.4 Å resolution (Bouyain *et al*., 2005)) for the tethered conformation and PDB:3U7U (3.03 Å (Liu *et al*., 2012)) for the active conformation. The E452K mutation was analyzed using PDB:3U7U, the extracellular structure of an active ERBB4 dimer with bound NRG-1. The L798R mutation was analyzed using PDB:3BCE, the intracellular structure of an active ERBB4 asymmetric kinase dimer and PDB:3BBT, the inactive form with bound active site inhibitor lapatinib (Qiu *et al*., 2008). The mutant rotamers were selected using the inbuilt tool of PyMOL (The PyMOL Molecular Graphics System, Version 2.4, Schrödinger, LLC.).

### Western blotting

Cells were lysed in lysis buffer containing 1% Triton X-100, 10 mM Tris-Cl, pH 7.4, 150 mM NaCl, 1 mM EDTA, 10 mM NaF and supplemented with Halt Protease and Phosphatase Inhibitor Cocktail (Thermo Scientific). Equal amounts of proteins were separated by SDS-PAGE, transferred to nitrocellulose membranes and analyzed using the following primary antibodies: anti-phospho-EGFR Tyr1086 (#2220), anti-EGFR (#2232, Cell Signaling Technology), anti-phospho-ERBB2 Tyr1248 (#2247; Cell Signaling Technology), anti-ERBB2 (MA5-14057; Invitrogen), anti-phospho-ERBB3 Tyr1289 (#4791; Cell Signaling Technology), anti-ERBB3 (#12708, Cell Signaling Technology), anti-phospho-ERBB4 antibodies: Tyr984 (#3790, Cell Signaling Technology), Tyr1162 (PAB0486, Abnova) and Tyr1284 (#4757, Cell Signaling Technology), anti-ERBB4 (clone E200, ab32375, Abcam), anti-phospho-AKT Ser473 (#4060; Cell Signaling Technology), anti-AKT (#2920; Cell Signaling Technology), anti-phospho-ERK Thr202/Tyr204 (#9101; Cell Signaling Technology), anti-ERK (#9102; Cell Signaling Technology) and anti-β-actin (A5441; Sigma Aldrich). Signals were detected using Odyssey CLx imaging system (LI-COR).

### Dimerization assays

To analyze ERBB4 homodimerization, COS-7 cells were seeded on 6-well plates and transiently transfected with FuGENE 6 and 1 *µ*g of the *pBABE-puro-gateway* constructs encoding wild-type or mutant ERBB4 or vector control. Next day, serum-free medium supplemented or not with 50 ng/ml NRG-1 was changed after washing the cells with PBS. After four-day serum starvation, cells were washed three times with ice-cold PBS and treated with 2 mM membrane impermeable BS_3_ crosslinker (Thermo Fisher) in PBS for 1 hour on ice. The reaction was quenched with ice-cold 50 mM Tris-HCl, pH 7.4, 150mM NaCl for 15 min on ice, after which the cells were washed three times with ice-cold PBS and lysed. The protein samples were run on 6 % SDS-PAGE gels and ERBB4 dimers were analyzed by western blotting.

To analyze ERBB4 heterodimerization, COS-7 cells were seeded on 10 cm dishes and transiently co-transfected with FuGENE 6 and 3 *µ*g + 3 *µ*g of the *pBABE-puro-gateway* constructs encoding ERBB4 variants or vector control with either EGFR, ERBB2 or ERBB3. Cells were serum starved for four days in the presence or absence of 50 ng/ml NRG-1 as described above and lysed. Lysates containing 500 *µ*g protein were precleared with 20 *µ*l protein G Sepharose beads (Cytiva) for 2 hours at +4 °C and subjected to immunoprecipitation by overnight incubation with either anti-ERBB4 (HFR-1 or E200), anti-EGFR, anti-ERBB2 or anti-ERBB3 antibodies and subsequent 2-hour incubation with 20 *µ*l protein G Sepharose beads. Beads were washed four times with 1 ml lysis buffer and boiled in SDS-PAGE loading buffer. Co-immunoprecipitating ERBB receptors were analyzed by western blotting.

### Patient data

For the analysis of gene alteration frequencies, cancer type and co-occurring alterations in ERBB4-mutant cancer patients, clinical cancer sample data were obtained from cBioPortal (https://cbioportal.org), AACR GENIE (https://genie.cbioportal.org), COSMIC (https://cancer.sanger.ac.uk/cosmic) in January 2024.

Neratinib efficacy data of patients harboring *ERBB4* alteration, enrolled in PUMA-NER-5201, the SUMMIT trial (NCT01953926), and treated with neratinib as a single agent (240 mg/day) were obtained from Puma Biotechnology and cBioPortal.

#### Drug sensitivity assays

IL3-independent Ba/F3 cells (20,000 cells/well) expressing ERBB4 variants were plated in 96-well plates in IL3-free medium containing or not 10 ng/ml NRG-1 and vector control cells were plated in medium containing IL3. The cells were incubated in the presence of 0.00067-2.5 *µ*M of neratinib (Puma Biotechnology), afatinib (Boehringer Ingelheim), dacomitinib (Cayman Chemicals) or DMSO only for 72 hours before measuring cell viability of quadruplicate samples with MTT assay (as described above). For PC-9 cells expressing ERBB4 variants, 3,000 cells/well were plated on 96-well plates. The next day, the growth medium was changed to a medium supplemented or not with 50 ng/ml NRG-1, and the cells were treated with 0.002 – 2.5 *µ*M osimertinib (SelleckChem) or DMSO in triplicate. Cell viability was measured with MTT assay after 72-hour incubation. The drugs and DMSO were dispensed with D300 Digital Dispenser (HP). GraphPad Prism 10 was used to fit dose-response curves with the variable slope equation and least squares regression and plotted by indicating mean ± standard deviation. For Ba/F3 experiments, the IC50 values were extrapolated from the fitted curves and normalized to that of wild-type ERBB4 of each independent experiment. For statistical analysis, normalized IC50 values of 2-3 independent experiments were analyzed by one-sample *t* test, adjusting the p-values for multiple comparisons by the Bonferroni method and considering *P* < 0.05 as significant.

### Long-term drug treatment assays

PC-9 cells expressing ERBB4 variants were plated on 12-well plates (25,000 cells/well). The following day, the growth medium was changed to a medium supplemented or not with 50 ng/ml NRG-1, and the cells were treated with 100 nM osimertinib (SelleckChem) or DMSO in triplicate. Medium was changed every 5 days. After 14-day incubation, the cells were washed with PBS, fixed with methanol (10 min) and stained with 0.5% crystal violet. After 15-minute incubation, the wells were washed three times with 2 ml of PBS, followed by one wash with 2 ml of ultrapure water (Milli-Q). After washes, the wells were allowed to air dry. The stained cells were imaged using Epson scanner, and the scanned images of three independent experiments were quantified in ImageJ, using the ColonyArea plugin (Schneider, Rasband and Eliceiri, 2012). One-way ANOVA was used to assess statistical significance, using Šídák’s multiple comparisons test.

### Mouse xenograft experiments

Seven-week-old female Rj:NMRI-Foxn1^nu/nu^ mice were purchased from Janvier Labs and were allowed to acclimate for at least six days prior to the initiation of the experiment at Turku Center for Disease Modeling (TCDM). Xenografts were established by subcutaneous injections of PC-9 cells expressing ERBB4 variants (5 x 10^6^ cells in 50% Matrigel matrix (#354234, BD Biosciences)) in the right flank of each mouse. Tumor growth was monitored one to three times a week by bilateral caliper measurements and the tumor volume was calculated using the formula V = 0.5 × length × width^2^. Tumors were allowed to reach 250 ± 50 mm^3^ in size on average before randomization into vehicle or osimertinib treatment groups (n > 4-5 or n > 11-13 mice per group, respectively). Powdered osimertinib (HY-15772, MedChemExpress) was added to normal chow mixture (V1554, ssniff Spezialdiäten) and pelleted by ssniff Spezialdiäten. The prepared chow containing 35 mg/kg osimertinib or not (vehicle) was fed to mice, and after 189 days, osimertinib treatment was withdrawn by changing the mice to receive normal chow. Mice were sacrificed at indicated time points according to our protocol approved by the Regional State Administrative of Southwestern Finland (license number: ESAVI/7740/2023). The animal study was carried out according to the regulations of the Finnish Act on Animal Experimentation (62/2006).

Tumor growth curves of individual mice and Kaplan-Meier curves of relapse-free survival were plotted with GraphPad Prism 10. Log rank test was used to compare the relapse-free survival of osimertinib-treated mice with tumors expressing either mutant or wild-type ERBB4.

### Single-cell RNA-sequencing analysis

Single-cell RNA sequencing data from PC-9 xenografts treated with vehicle or osimertinib (Kurppa *et al*., 2020) was downloaded from GEO (accession number GSE131604). Cellranger (version 7.1.0) aggregate analysis was performed on count data using default parameters to normalize count data by sequencing depth. The expression of ERBB4 ligands *NRG1*, *NRG2*, *NRG3, NRG4*, *HBEGF*, *EREG* and *BTC* was analyzed using Loupe Browser (version 8).

## RESULTS

### Selection of recurrent *ERBB4* mutations for functional characterization

To search for potentially actionable *ERBB4* mutations among the most recurrent cancer-associated *ERBB4* mutations, we conducted an *in silico* analysis of the curated set of non-redundant studies (n = 217) included in the cBioPortal, consisting of 70,655 patient samples from 147 cancer types (Supplementary Fig. S1A). *ERBB4* was altered in 3.5% of the analyzed samples in which missense mutations were the most common *ERBB4* alteration (87% of *ERBB4* altered samples) (Fig. 1A). The analysis identified 2,117 somatic *ERBB4* missense mutations of which 182 (9%) were annotated as putative drivers in cBioPortal based on OncoKB curated evidence (Chakravarty *et al*., 2017; Suehnholz *et al*., 2024) (Fig. 1A). *ERBB4* missense mutations were found to be most frequent in skin cancers (30% in non-melanoma, 14% in melanoma) and least frequent in embryonal (0.2%) cancers (Supplementary Fig. S1B).

**Figure 1.**
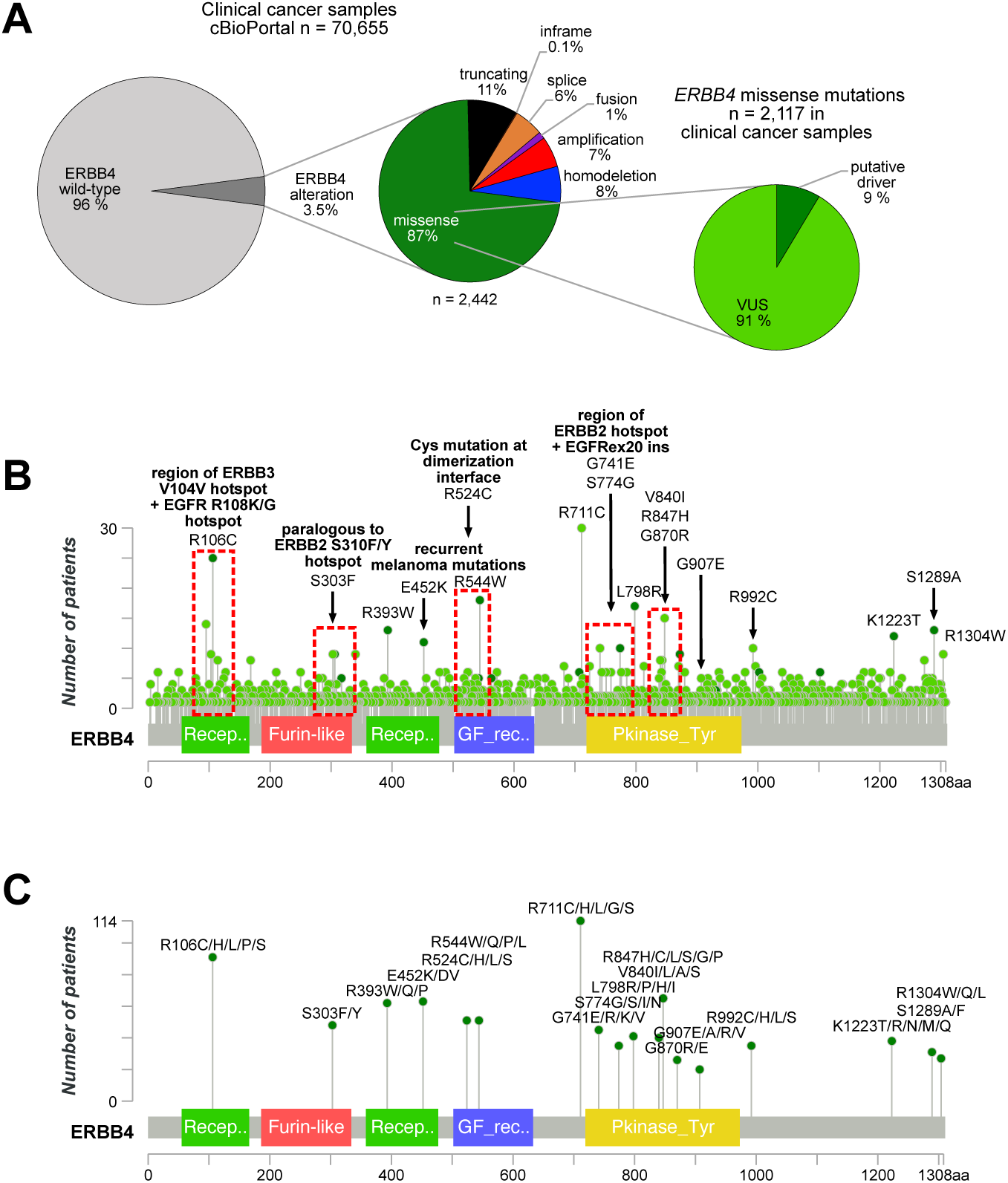
Cancer-associated *ERBB4* mutations selected for functional characterization. **A)** Left and middle: fractions of *ERBB4* alterations in clinical cancer samples reported in cBioPortal (cbioportal.org) curated non-redundant studies (January 2024). Right: fractions of putative driver cases and variant of unknown significance (VUS) cases in all *ERBB4* missense mutated cancer samples (based on OncoKB curated evidence on putative oncogenicity of a given alteration). Of the *ERBB4* missense mutations reported in cBioPortal (n = 2,117), 9% were annotated as putative drivers, and 91% as variants of unknown significance (VUS). **B)** Lollipop diagram depicting the number of missense mutations at the indicated ERBB4 amino acid residues reported in cBioPortal. Light green denotes VUS and dark green denotes putative driver mutation based on OncoKB annotations. Regions of interest (red rectangles) demonstrate 1) paralogous mutations to known activating mutations described for other oncogenic ERBB family members and/or 2) location at dimerization interface suggesting functional relevance. The 18 recurrent missense mutations chosen for analyses are indicated. **C)** Lollipop diagram showing the number of patients with the selected *ERBB4* mutations, or less frequent mutations targeting the same amino acid, reported in cBioPortal, AACR GENIE (genie.cbioportal.org) and COSMIC (cancer.sanger.ac.uk) (redundant cases removed). The MutationMapper tool of cBioPortal was used to visualize the data.

While the missense mutations were distributed across the 1,308 amino acid sequence of ERBB4, clusters of mutations were identified in specific regions that are i) paralogous to activating mutations described for other oncogenic ERBB family members and/or ii) around ERBB4 dimerization interfaces, suggesting functional relevance (red boxes in Fig. 1B). We selected in total 18 *ERBB4* missense mutations (indicated in Fig. 1B) that were recurrent and located in the abovementioned regions of interest for functional characterization (indicated in Fig. 1B and Supplementary Fig. S1C) – hypothesizing that these mutations would be actionable. Of the different mutants at the same position of ERBB4 amino acid sequence, the most recurrent amino acid change was selected for characterization.

To more comprehensively assess the recurrence of the selected *ERBB4* missense mutations (and the less frequent mutations at the same amino acid residue) in cancer patients, all such patient cases listed in three publicly available cancer registries (cBioPortal, AACR GENIE and COSMIC) were combined and the non-overlapping cases are shown in Figure 1C. The numbers of patients ranged from 114 (R711/H/L/G/S mutations) to 20 (G907E/A/R/V mutations) (Fig. 1C).

### Majority of the recurrent ERBB4 mutations are transforming in Ba/F3 or MCF10a cells

To study the functional significance of the 18 selected recurrent *ERBB4* mutations, we set out to screen their ability to transform (as outlined in Fig. 2A) two different non-tumorigenic cell models: murine lymphoid Ba/F3 cells and human mammary epithelial MCF10a cells. The ERBB4 JM-a CYT-2 isoform was used in the studies based on previous findings suggesting that JM-a CYT-2 is the more oncogenic ERBB4 isoform of the cancer-associated isoforms (Veikkolainen *et al*., 2011) in hematopoietic cell contexts (relevant for the Ba/F3 cell model) (Määttä *et al*., 2006; Chakroborty *et al*., 2022) and in numerous mammary epithelial cell contexts (relevant for the MCF10a cell model) (Sartor *et al*., 2001; Junttila *et al*., 2005; Vidal *et al*., 2007; Muraoka-Cook *et al*., 2009). Both the cell lines were retrovirally transduced to stably express the selected ERBB4-mutants, wild-type ERBB4 or eGFP as a vector control. To assess the transforming potential of the ERBB4-mutants, cell proliferation was measured upon depriving Ba/F3 cells of exogenous IL3 or MCF10a cells of EGF, of which the proliferation of these cell lines is normally dependent on. The proliferation assays were conducted in the presence or absence of the ERBB4 ligand NRG-1. Eleven out of the 18 mutants were identified as significantly more transforming than wild-type ERBB4 in either of the two cell models, in the presence or absence of NRG-1, as summarized in a heatmap (Fig. 2B). Three of these mutants (S303F, E452K and L798R) demonstrated enhanced transforming activity in both cell models, implying context-independent transforming potential. Of note, wild-type ERBB4 was also capable of transforming both Ba/F3 and MCF10a cells in the presence but not in the absence of NRG-1 (Fig. 2C-D).

**Figure 2.**
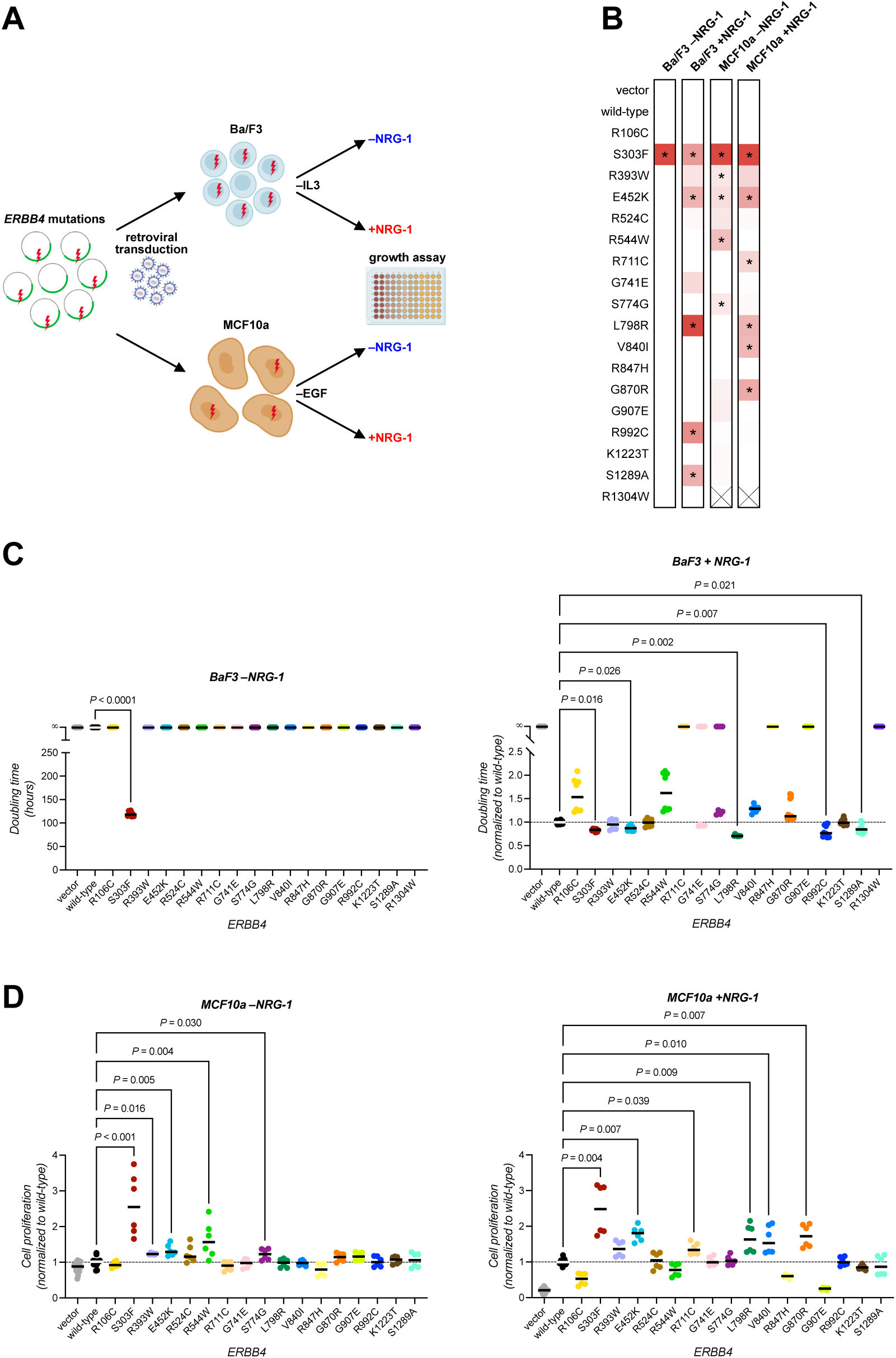
Recurrent *ERBB4* mutations have transforming potential. **A)** Outline of the screen used to assess the transforming potential of 18 recurrent *ERBB4* mutations in non-tumorigenic Ba/F3 murine lymphoid cells and in MCF10a human mammary epithelial cells. Image created with BioRender.com. **B)** Summary of the transforming activity of *ERBB4* mutations compared to wild-type *ERBB4*. Heatmap summarizes the data from C and D (*, *q* < 0.05; Kruskal Wallis test. P-values were controlled for false discovery rate by two-stage linear step-up method of Benjamini, Krieger and Yekutieli). X indicates a cell line that could not be generated despite several attempts. **C)** The effect of ERBB4 variants on Ba/F3 cell doubling time upon IL3 deprivation in the presence or absence of 20 ng/ml NRG-1. Cell proliferation was analyzed with MTT assay over time, and doubling times were calculated (Methods). Doubling times in the presence of NRG-1 were normalized to the average of wild-type ERBB4 expressing cells of each independent experiment. **D)** The effect of ERBB4 variants on MCF10a cell growth upon EGF deprivation in the presence or absence of 50 ng/ml NRG-1. Cell proliferation was analyzed with the MTT assay on day 0 and 8 and fold changes in cell proliferation were calculated and normalized to the average of wild-type ERBB4 expressing cells. The dashed horizontal lines in C and D indicate the average of cells expressing wild-type ERBB4.

Remarkably, ERBB4 S303F was the only variant able to promote IL3-independent growth of Ba/F3 cells without NRG-1 (Fig. 2C). Five out of the 18 mutants (S303F, E452K, L798R, R992C and S1289A) promoted IL3-independent Ba/F3 cell proliferation in the presence of NRG-1 more potently than wild-type ERBB4 (*q* < 0.05; Fig. 2C). In MCF10a cells, five mutants (S303F, R393W, E452K, R544W and S774G) were more potent in promoting EGF-independent cell proliferation in the absence of NRG-1 than wild-type ERBB4 (*q* < 0.05; Fig. 2D). In the presence of NRG-1, six not completely overlapping mutants (S303F, E452K, R711C, L798R, V840I and G870R) promoted MCF10a cell proliferation better than wild-type ERBB4 (*q* < 0.05; Fig. 2D).

Taken together, these analyses indicate a potential oncogenic role for 11 recurrent ERBB4 mutations. Three mutants (S303F, E452K and L798R) were strongly transforming with the ability to transform both cell models, S303F being unique in its ability to transform both models in the absence of NRG-1.

### Structural analysis of the transforming ERBB4 mutations

To study the mechanisms underlying the transforming potential of the three most potent ERBB4-mutants, their potential structural impact was assessed. The S303 residue of ERBB4 is located at the extracellular β-helical domain II (Fig. 3A) (in the sixth module out of seven small disulfide-containing modules (Cho and Leahy, 2002)). In a monomeric structure of ERBB4 in the inactive tethered conformation, the side chain of S303 faces solvent and is hydrogen bonded to the side chain of T284 of module 5, whereas in a dimeric form the residue is in close proximity of the dimerization arm of the neighboring subunit (Fig. 3B). Two features of the S303F mutation would likely contribute to enhanced receptor activity. Firstly, while the side chain of S303 is hydrophilic and faces the solvent in the tethered structure, the phenylalanine residue in the S303F-mutant prefers a hydrophobic environment. Secondly, in the active dimer S303F would be located within a hydrophobic pocket interacting with the side chains of L275 and Y268 (π−π stacking). This would likely result in ERBB4 S303F favoring the active, open conformation over the inactive tethered conformation, and stabilizing interactions in the active dimer. The paralogous mutation S310F in ERBB2 was shown to stabilize a complex with ERBB3 and NRG-1 (Diwanji *et al*, 2021).

**Figure 3.**
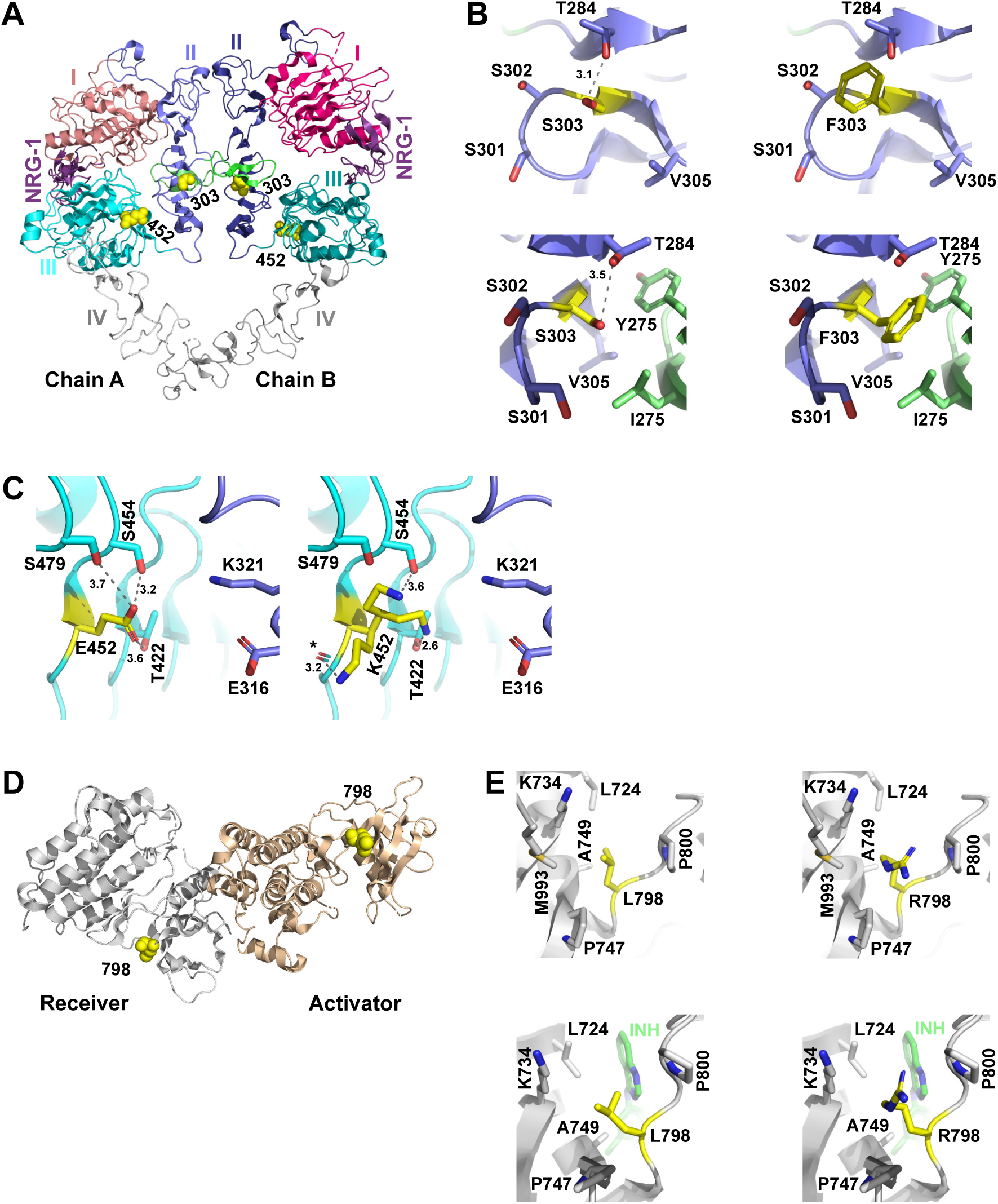
Structural analysis of the S303F, E452K and L798R mutations. **A)** Structure of the ERBB4 extracellular domains in an NRG-1 -bound active homodimer (PDB:3U7U). Yellow spheres indicate the location of the S303F and E452K mutations. **B)** The S303F mutation site, shown in the presence of S303 (left) or F303 (right). The upper and lower panels represent the tethered (PDB: 2AHX) and the active (PDB: 3U7U) conformations, respectively. **C)** The E452K mutation site in the active dimer structure (PDB:3U7U), shown in the presence of E452 (left) or K452 (right). Three potential lysine rotamers are shown. **D)** The structure of the active ERBB4 asymmetric kinase dimer (PDB:3BCE) with yellow spheres indicating the location of L798 in both the activator and receiver kinases. **E)** The L798R mutation site, shown in the presence of L798 (left) or R798 (right). The upper and lower panels represent the active kinase in the asymmetric dimer (PDB: 3BCE) and the inactive kinase [PDB: 3BBT, inactive conformation with bound lapatinib (INH)], respectively.

The E452K mutation is also extracellular and located at the interface of domains II and III (Fig. 3A). However, no direct contacts can be seen between these domains; a significant change in conformation would be needed for interdomain interactions, e.g. with E316 and/or K321 of domain II (Fig. 3C). E452 seems to form a stable hydrogen-bonding network with the surrounding residues T422, S454 and S479, whereas in the mutant the lysine residue has multiple options to form hydrogen bonds, which may affect the dynamics of the interface and dimer stability. In the tethered structure, E452 is facing the solvent and the mutation E452K is unlikely to cause any major effect.

Leucine 798 is positioned at the active site in the kinase domain (Fig. 3D) and within van der Waals interaction distance to the inhibitor (lapatinib) in the 3BBT structure (Fig. 3E, *lower panels*). In the active kinase domain structure of EGFR with bound ATP analog (PDB:2GS6), an equivalent interaction between the paralogous leucine residue and an ATP analog exists. In the L798R-mutant, the bulky arginine residue may affect substrate binding at the active site and catalytic activity; e.g. R798 could form π−π stacking interactions with the adenine ring of ATP leading to increased activity but this would require local conformational changes at the active site.

### Transforming ERBB4 mutations are activating and co-operate with other ERBB family members

To test assess whether the three most potent ERBB4-mutants were transforming due to gain-of-function effect as proposed by the structural analyses, ERBB4 phosphorylation levels and downstream signaling pathway activation were analyzed in MCF10a cells (Fig. 4A). MCF10a cells express endogenously other ERBB family receptors, allowing heterodimerization with ERBB4, and thus, making MCF10a a relevant cellular context to study mutant ERBB4 signal transduction (Fig. 4A). Upon eight-day EGF-deprivation, corresponding to the conditions of the transformation assay (Fig. 2D), the strongly transforming ERBB4-mutants had indeed greater autophosphorylation levels both in the presence and absence of NRG-1. Additionally, EGFR, ERBB2, and ERBB3 were more phosphorylated in mutant ERBB4-expressing cells than in wild-type ERBB4-expressing or vector control cells in the presence and absence of NRG-1, most prominently in ERBB4 S303F-expressing cells (Fig. 4A, *lanes 5 and 6*). Consistently, the ERBB4 downstream signaling pathways were also more active in cells expressing ERBB4-mutants. In summary, these data indicate that S303F, E452K and L798R are activating, gain-of-function ERBB4 mutations that may co-operate with other ERBB receptors in malignant transformation.

**Figure 4.**
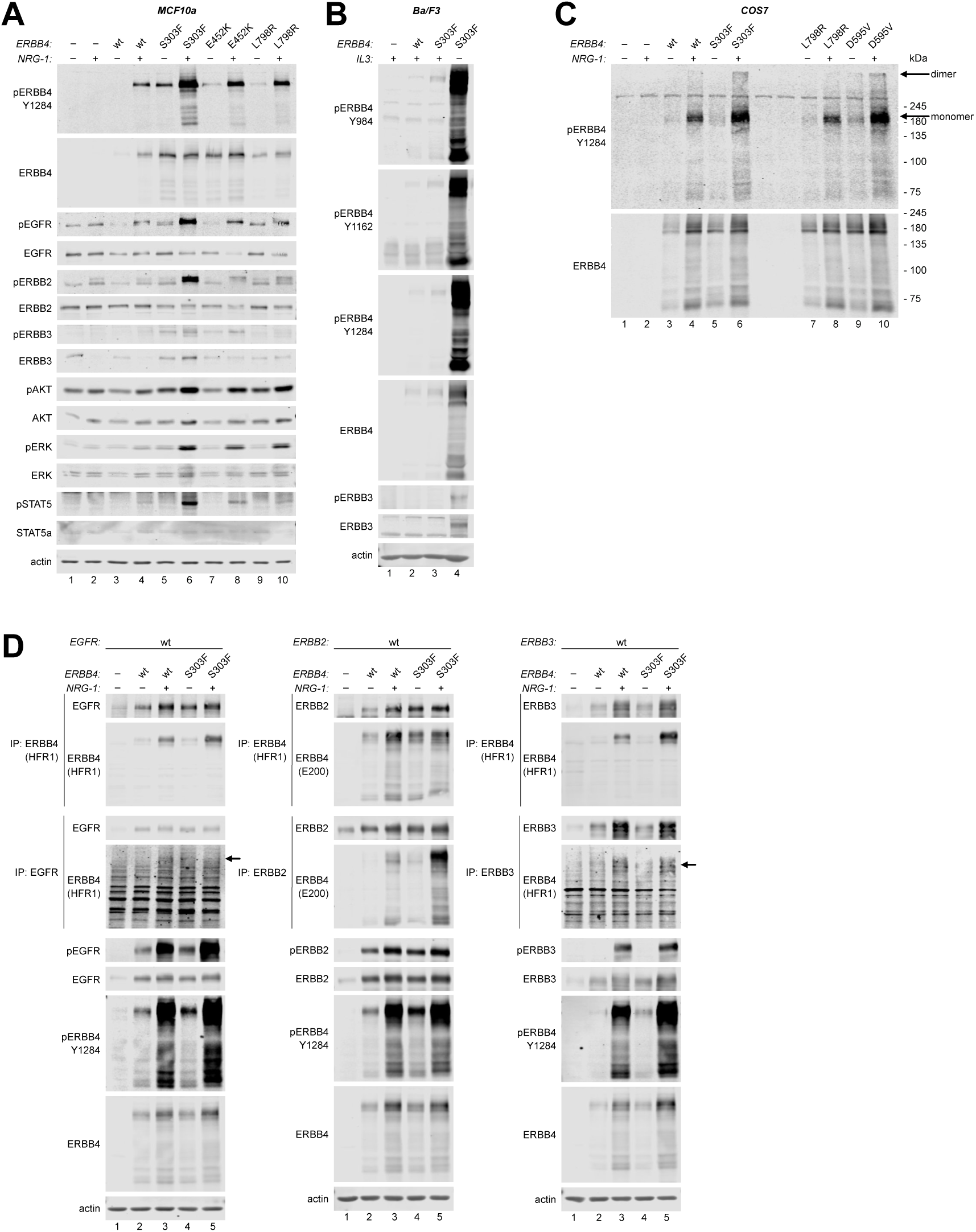
Strongly transforming ERBB4 mutations are activating and co-operate with other ERBB receptors. **A)** MCF10a cells stably expressing ERBB4 variants or vector control (-) were cultured in the absence of EGF and in the absence or presence of 50 ng/ml NRG-1 for eight days. Lysates were analyzed by western blot using beta-actin as a loading control. **B)** Ba/F3 cells stably expressing the indicated ERBB4 variants or vector control were cultured in the presence or absence of IL3 and analyzed by western blot. **C)** COS7 cells were transfected with constructs encoding ERBB4 variants or vector control and cultured in the absence of serum and in the presence or absence of 50 ng/ml NRG-1 for four days. Cells were incubated with membrane impermeable BS_3_ crosslinker and analyzed for phosphorylated ERBB4 dimers by western blot. **D)** COS7 cells co-transfected with an ERBB4 variant or vector control together with either EGFR, ERBB2 or ERBB3 were cultured as in C. Cells were lysed and analyzed for heterodimerization by immunoprecipitation (IP) using either anti-ERBB4, anti-EGFR, anti-ERBB2 or anti-ERBB3 antibodies. IP samples and total lysates were analyzed by western blot using the indicated antibodies.

In contrast, when MCF10a cells were not subjected to eight-day EGF-deprivation but instead to short-term overnight serum starvation followed by NRG-1 stimulation for 10 minutes, there was no difference in ERBB4 phosphorylation between the transforming mutants and wild-type ERBB4 (Supplementary Fig. S2A). Additionally, a gradual loss of wild-type ERBB4 but not ERBB4 S303F expression was observed over prolonged culture (≥10 weeks) of the stable MCF10a transductants (Supplementary Fig. S2B-C) despite constant antibiotic selection to maintain the ERBB4-expressing pool of cells. The loss of wild-type ERBB4 occurred while maintaining the cells in the presence of EGF (Supplementary Fig. S2B), suggesting that wild-type ERBB4 JM-a CYT-2, but not the S303F-mutant, provides selection disadvantage in the absence of transformation pressure. However, the EGF-independent growth of MCF10a cells mediated even by wild-type ERBB4 (Fig. 2D), together with the increased activation of the strongly transforming mutants observed upon EGF-deprivation (Fig. 4A) imply that ERBB4 activation favors the survival of breast epithelial cells upon deprived conditions providing selection pressure for transformed cells.

Since all three strongly transforming ERBB4-mutants, particularly ERBB4 S303F, promoted activation of all ERBB receptors in MCF10a cells (Fig. 4A), we hypothesized that ERBB4 heterodimerization might also be associated with cell transformation in Ba/F3 cells. Ba/F3 cells are known to endogenously express low levels of ERBB3 but no other ERBB receptors (Riese *et al*., 1995), and up-regulation of endogenous ERBB3 expression is associated with Ba/F3 cell transformation promoted by other ERBB receptors, such as oncogenic mutants of EGFR (Supplementary Fig. S3A). Indeed, ERBB4 S303F-transformed IL3-independent Ba/F3 cells had highly increased expression and phosphorylation levels of both ERBB4 and ERBB3 compared to non-transformed cells growing with IL3 (Fig. 4B). While ERBB4 S303F was more phosphorylated than wild-type ERBB4 at three different tyrosine residues (Y984, Y1162, Y1284) even in the presence of IL3, ERBB3 activation was only detectable in transformed, IL3-independent Ba/F3 cells.

To further investigate the role of ERBB3 upregulation in ERBB4-mediated Ba/F3 cell transformation, ERBB3 was knocked down in vector control cells and cells expressing ERBB4 wild-type or S303F using doxycycline-inducible murine *Erbb3* targeting shRNA (Supplementary Fig. S3B). IL3-deprivation in the presence or absence of doxycycline revealed that *Erbb3* knockdown significantly impaired ERBB4 S303F-mediated, IL3-independent growth of Ba/F3 cells in the absence of NRG-1 (Supplementary Fig. S3C-D, *P* < 0.0001). In contrast, *Erbb3* knockdown did not inhibit the growth of IL3-dependent cells, nor cells growing in the presence of NRG-1. Together, these data suggest that while ERBB4 can be transforming in the absence of other ERBB receptors, mutant ERBB4 co-operates with ERBB3 to promote ligand-independent cell transformation.

### ERBB4 S303F stabilizes dimers

Considering the ability of ERBB4 S303F to broadly activate other ERBB receptors, and the structural analyses suggesting enhanced dimerization ability for ERBB4 S303F, we assessed the capability of ERBB4 S303F to form homodimers and heterodimers with other ERBB receptors. These experiments were performed with COS7 cells, which are devoid of endogenous ERBB4 and have low endogenous EGFR, ERBB2 and ERBB3 levels (Sawano *et al*., 2002). Although ERBB4 S303F-mutant was slightly more phosphorylated after overnight serum starvation and 10-minute NRG-1 stimulation than wild-type ERBB4 in COS7 cells (Supplementary Fig. S4A, *lanes 3 and 5*), the difference in ERBB4 phosphorylation between S303F and wild-type amplified over time in serum starvation (Supplementary Fig. S4B). This is consistent with the observations of more prominently elevated phosphorylation levels of mutant ERBB4 compared to wild-type ERBB4 upon prolonged growth signal deprivation in both MCF10a (upon EGF deprivation: Fig. 4A *vs* Supplementary Fig. S2A) and Ba/F3 cells (upon IL3 deprivation Fig. 4B, *lanes 3 vs 4*).

We selected four-day serum starvation for further experiments, demonstrating robust difference in ERBB4 wild-type and S303F phosphorylation levels, similar to what was observed in MCF10a and Ba/F3 cells. ERBB4 homodimers were assessed by crosslinking cell surface proteins with a cell membrane impermeable BS_3_, enabling detection of ERBB4 dimers as high molecular weight species of ERBB4 in western blot. Another activating extracellular ERBB4 mutation, D595V, was used as a positive control, as we have previously demonstrated D595V to stabilize ERBB4 dimers using the same assay (Kurppa *et al*., 2016). As predicted by the structural analyses, S303F resulted in more abundant active, phosphorylated ERBB4 dimers than wild-type ERBB4 in the presence of NRG-1, while the activating intracellular domain mutation L798R, that served as a negative control for dimer stabilization, did not (Fig. 4C).

To study the capability of ERBB4 S303F to form ERBB heterodimers, wild-type EGFR, ERBB2 or ERBB3 was co-transfected with ERBB4 wild-type or S303F, and heterodimerization was analyzed by co-immunoprecipitation. Interestingly, the expression levels of transiently transfected ERBB4 and ERBB3 were greater in the cells serum-starved for four days in the presence of NRG-1 than in its absence (Fig. 4D), suggesting that high expression of ERBB4 and ERBB3 provides growth-advantage for cells under serum starvation. Taking into account these expression level differences, ERBB4 S303F did indeed co-immunoprecipitate more efficiently than wild-type ERBB4 with ERBB2 and EGFR both in the presence or absence of NRG-1 (Fig. 4D), demonstrating that the S303F mutation promotes the formation of ERBB heterodimers. Surprisingly, no difference in co-immunoprecipitation with ERBB3 was seen between S303F-mutant and wild-type ERBB4, although the experiments in Ba/F3 cells (Fig. 4B) suggested functional role for ERBB3 heterodimerization upon ERBB4 S303F-mediated establishment of IL3-independence. However, the COS7 cells expressing the S303F-mutant demonstrated greater phosphorylation levels of both ERBB4 and ERBB3 as compared to the cells expressing wild-type ERBB4, suggesting that ERBB4 S303F together with ERBB3 promoted cell survival during the four-day serum starvation.

Taken together, this *in vitro* evidence is consistent with the structural predictions that ERBB4 S303F can mediate its activating functions by stabilizing homo- and heterodimers with other ERBB receptors but also that the heterodimerization is likely cell/tissue context-dependent.

### Clinical efficacy of single-agent pan-ERBB inhibitor neratinib in patients with ERBB4 mutations

Despite the recurrence of *ERBB4* mutations in cancer and the existence of clinically used pan-ERBB inhibitors that potently also block ERBB4 activity, there is very limited data about the clinical actionability of *ERBB4* mutations. However, the SUMMIT basket trial (clinicaltrials.gov identifier NCT01953926) assessing the efficacy of pan-ERBB inhibitor neratinib in patients with somatic *ERBB2*, *ERBB3* or *ERBB4* mutations, included seven patients harboring *ERBB4* alterations. Four of the seven patients had an *ERBB4* mutation characterized in this study. Two of the three patients that were qualified for the SUMMIT trial due to a mutation in *ERBB4*, with no other qualifying mutations in ERBB family genes, had an *ERBB4* mutation characterized in this study to be transforming (R544W and V840I) (Table 1). Yet, neither of these patients, nor the patient with an ERBB4 VUS N465K, responded to neratinib, progressive disease being the best response (Table 1).

**Table 1.** Patients harboring *ERBB4* alterations and treated with neratinib in the SUMMIT trial. *ERBB4* alteration harboring patients enrolled in PUMA-NER-5201, the SUMMIT trial (NCT01953926), based on a mutation in *ERBB4* (A) or in *ERBB2*(/*ERBB3*) (B) were treated with neratinib as a single agent (240mg/day). Outcomes of neratinib treatment, detailed cancer type and co-altered genes are presented. Patients whose tumors harbored *ERBB4* mutations functionally characterized in this work are indicated with red rectangles. PFS, progression-free survival; OS, overall survival.

| Qualifying mutation for SUMMIT trial in *ERBB4* |  |  |  |  |  |  |  |
| --- | --- | --- | --- | --- | --- | --- | --- |
| Qualifying mutation |  | Co-altered genes | Cancer type | Prior lines of therapy | Best response | PFS (months) | OS (months) |
| *ERBB4* N465K |  | PIK3CA | ovarian clear cell carcinoma | 4 | PD | 1.1 | 2.2 |
| *ERBB4* R544W |  | TP53 | rectal adenocarcinoma | 6 | PD | 1.7 | 14.3 |
| *ERBB4* V840I |  | TP53, HER2 amp, CCNE1 amp | peri-ampullary adenocarcinoma | 1 | PD | 1.7 | not evaluable (withdrew consent) |
| Qualifying mutation for SUMMIT trial in another *ERBB* gene |  |  |  |  |  |  |  |
| Qualifying mutation | Co-occurring *ERBB4* mutation | Co-occurring putative drivers | Cancer type |  | Best response | PFS (months) |  |
| *ERBB2* S310F | *ERBB4* Q132* | *ERBB2* (D277H), KDM6A, ARID1A, TERT, ERCC4, CDKN2A del, CDKN2B del | bladder urothelial carcinoma |  | SD | 3.5 |  |
| *ERBB2* S310F | *ERBB4* R711C | TP53, CDKN2A, TERT, TGBR1 | cutaneous melanoma |  | SD | 1.3 |  |
| *ERBB2* D769Y | *ERBB4* L798R, N138S | *ERBB2* amp, TP53, APC, ARID1B, MCL1 amp, MYC amp, CCND3 amp, CCND1 amp, FGF19 amp, FGF4 amp, FGF3 amp, VEGFA amp, RECQL4 del | esophageal adenocarcinoma |  | PD | 1.9 |  |
| *ERBB2* S310Y | *ERBB4* amp | TP53, TERT, ARID1A, EPHA7, RB1, AXL amp, MCL1 amp, AKT2 amp, MYCL amp | bladder/urinary tract cancer |  | PD | 1.2 |  |

Additionally, four patients that were enrolled to SUMMIT due to a mutation in *ERBB2* had co-occurring *ERBB4* mutations or *ERBB4* amplification (Table 1), including two mutations characterized as transforming in this study: the strongly transforming L798R mutation and R711C that was transforming in MCF10a cells but not in Ba/F3 cells. The ERBB4 L798R-mutant patient harbored also another ERBB4 VUS, N138S, and did not respond to neratinib while the patient harboring ERBB4 R711C had stable disease as the best response. It is noteworthy, that all the four patients whose *ERBB4* mutation is transforming *in vitro* had a co-occurring *TP53* alteration, and *TP53* mutations were associated with lack of response to neratinib in the SUMMIT trial (Hyman *et al*., 2018).

### Activating ERBB4 mutations are sensitive to clinically available pan-ERBB inhibitors

Next, we set out to determine the sensitivity of the strongly transforming ERBB4-mutants (S303F, E452K, L798R) to neratinib and the two other clinically used second-generation pan-ERBB inhibitors that are known to inhibit ERBB4 at a low nanomolar range, afatinib and dacomitinib (Davis *et al*., 2011). All the three ERBB4 mutations rendered NRG-1-dependent Ba/F3 cells similarly sensitive to neratinib, afatinib and dacomitinib as wild-type ERBB4 (Fig. 5A-B; *P* > 0.05). Although the patient harboring ERBB4 L798R mutation did not respond to neratinib in the SUMMIT trial (Table 1), and L798R is located in the ATP-binding pocket where it could potentially interfere with the binding of TKIs (Fig. 3D-E), these *in vitro* data showed that L798R did not mediate resistance to neratinib, afatinib or dacomitinib.

**Figure 5.**
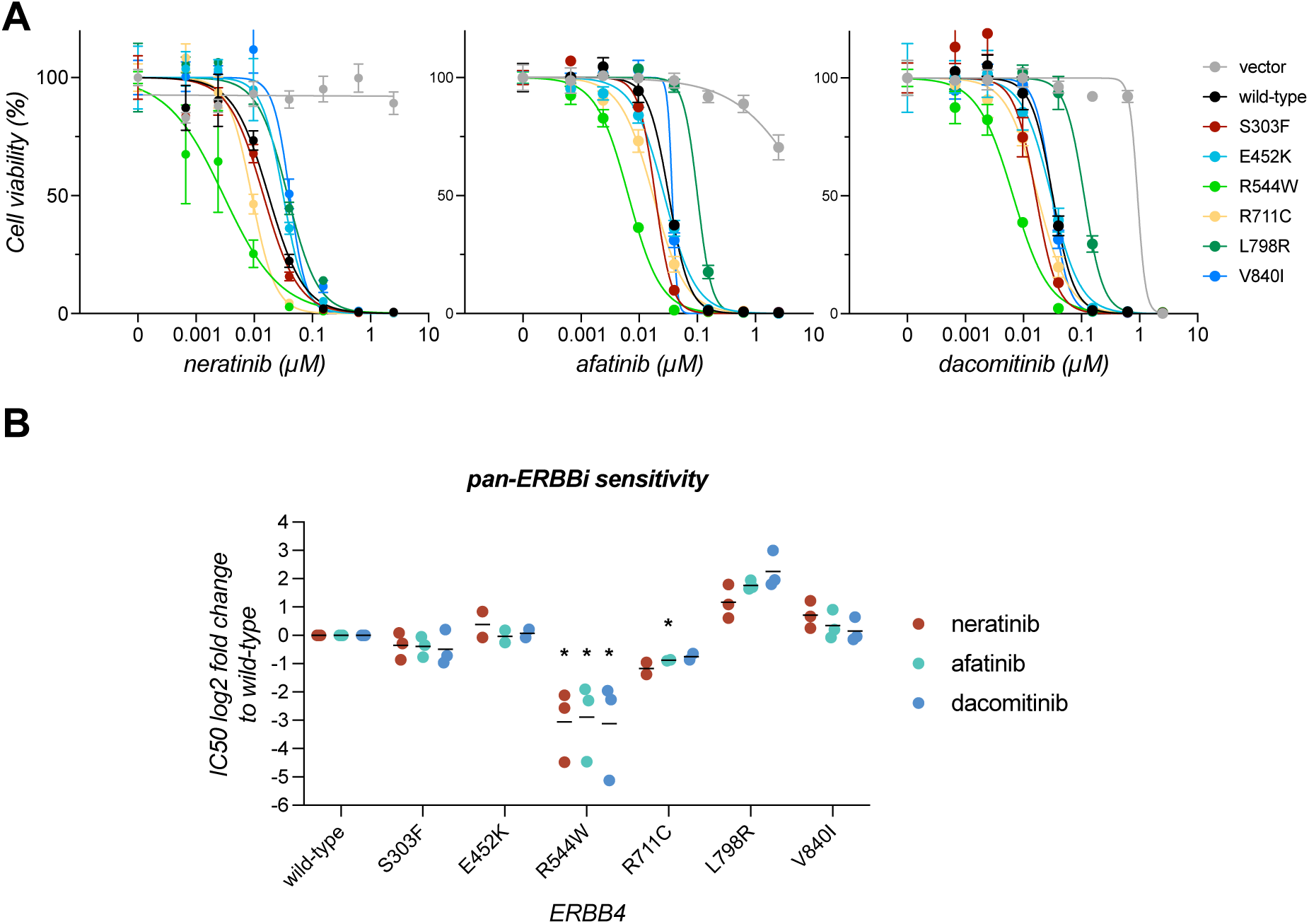
Transforming ERBB4 mutants are sensitive to pan-ERBB inhibitors. **A)** Sensitivity of Ba/F3 cells expressing different ERBB4 variants to pan-ERBB inhibitors. Ba/F3 cells expressing ERBB4 variants, growing in the absence of IL3 and presence of NRG-1, and vector control cells growing in the presence of IL3 were treated with the indicated concentrations of pan-ERBB inhibitors (neratinib, afatinib or dacomitinib) for 72 hours and cell viability was measured with the MTT assay. A representative dose response curve with mean and SD of quadruplicate analyses of one of 2-3 independent experiments is shown for each inhibitor. **B)** Fold changes in IC50 for Ba/F3 cells expressing ERBB4 variants, compared to Ba/F3 cells expressing wild-type ERBB4 are shown for each replicate experiment from A. *, *P* < 0.05; one-sample *t* test, p-values corrected for multiple comparisons by Bonferroni method. Horizontal lines indicate geometric means of the replicate experiments.

In addition, we analyzed the pan-ERBB inhibitor sensitivity of the three other transforming ERBB4-mutants detected in the SUMMIT trial patients who did not respond to neratinib (R544W, R711C, V840I). Cells expressing ERBB4 V840I were equally sensitive with wild-type ERBB4 to all the three pan-ERBB inhibitors (Fig. 5A-B; *P* > 0.05), whereas cells expressing the R544W-mutant were significantly more sensitive (neratinib *P* = 0.025, afatinib *P* = 0.038, and dacomitinib *P* = 0.041). Also, the cells expressing ERBB4 R711C trended towards being more sensitive to pan-ERBB inhibition than the cells expressing wild-type ERBB4 but the comparison reached statistical significance only with afatinib (*P* = 0.047). These *in vitro* findings demonstrate that the unresponsiveness of the patients harboring ERBB4 R544W, V840I, R711C and L798R mutations to neratinib in the SUMMIT trial cannot be attributed to these being *bona fide* resistance mutations to neratinib. Instead, these data suggest that all the analyzed transforming ERBB4 mutations are targetable with clinically available second-generation pan-ERBB inhibitors.

### Characteristics of patient tumors harboring transforming ERBB4 mutations

Analyses of the patient cases included in cBioPortal, AACR GENIE and COSMIC showed that all the three mutants were highly cancer type specific, S303F occurring mostly in breast cancer (69%, 25/36), E452K in melanoma (85%, 52/61), and L798R in cancers of the gastrointestinal tract (esophagogastric 47%, 15/32 and colorectal 32%, 10/32) (Supplementary Fig. S5A-B). Next, we investigated the frequency of co-occurring ERBB alterations (Supplementary Fig. S5C-D), considering the potential co-operation of heterodimers and recent studies showing that ERBB genes often harbor multiple weaker mutations *in cis*, i.e. compound mutations, resulting in enhanced receptor activation and oncogenicity (Saito *et al*., 2020; Koivu *et al*., 2024). Notably, S303F mutation occurred as the single ERBB alteration in 84% of the cases, and thus, the patients had significantly less co-occurring ERBB alterations than patients with any *ERBB4* alteration (*P* = 0.037), consistent with our observations that S303F is a strongly activating mutation. In contrast, E452K had significantly more co-occurring ERBB alterations (*P* = 0.0005), of which the majority were ERBB4 compound mutations, potentially indicative of co-operation of multiple weaker ERBB4 mutations. L798R, in turn, occurred as a single ERBB alteration in 65% of the cases, similar to clinical cancer samples with any *ERBB4* (64%)*, EGFR* (62%) *ERBB2* (70%), or *ERBB3* (64%) alteration (Supplementary Fig. S5D).

Taken together, these clinical sample data are in line with our *in vitro* results, suggesting that S303F is a potential driver mutation in breast cancer.

### Activating ERBB4 mutations drive resistance to EGFR-targeted therapy in EGFR-mutant NSCLC cells

Intriguingly, the *ERBB4* mutations identified in patients who developed resistance to EGFR-targeted therapy (Cremolini *et al*., 2019; Jänne *et al*., 2022), include the same mutation or mutation in the same residue as analyzed in the current study: the strongly transforming S303F or L798I. This points to the possibility that mutant ERBB4 could promote resistance to targeted therapies. To address this, the effect of activating ERBB4 mutations on EGFR inhibitor response in EGFR-mutant NSCLC cells was assessed. For this, we transduced PC-9 cells, lacking endogenous ERBB4, with wild-type ERBB4 or four activating ERBB4-mutants: S303F and L798I identified in this study, as well as previously identified E715K (Chakroborty *et al*., 2022) and K935I (Kurppa *et al*., 2016). Consistent with previous experiments (Fig. 4, (Kurppa *et al*., 2016; Chakroborty *et al*., 2022)), the strongly activating S303F and E715K-mutants demonstrated increased ERBB4 activation compared to wild-type ERBB4 following short-term stimulation with NRG-1 (Fig. 6A). On the other hand, the L798R and K935I-mutants did not exhibit enhanced ERBB4 activation, suggesting some context-dependency in mutant ERBB4 activation, at least in the steady-state setting. In short-term, three-day experiments, the mutant ERBB4-expressing PC-9 cells did not demonstrate any change in response to EGFR inhibitor osimertinib, either in the presence or absence of NRG-1 (Fig. 6B), when compared to wild-type ERBB4-expressing or vector control cells. It is noteworthy though that NRG-1 promoted a clear increase in osimertinib IC50 in all cell lines, indicating that NRG-1 (most likely through ERBB3) may affect EGFR inhibitor response.

**Figure 6.**
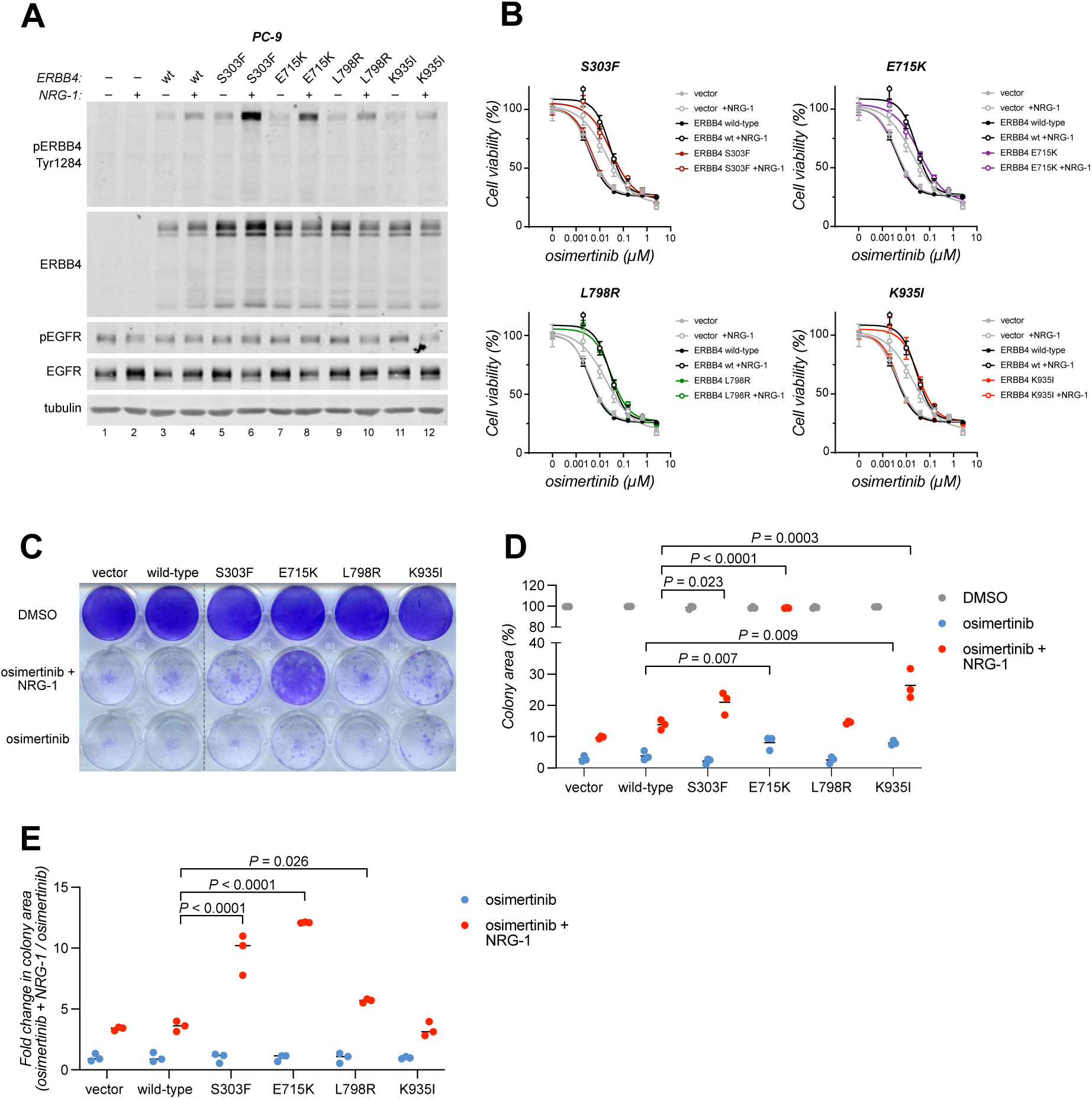
Activating ERBB4 mutations drive resistance to EGFR-targeted therapy in EGFR-mutant NSCLC cells. **A)** PC-9 cells stably expressing ERBB4 variants or vector control cells (-) were stimulated or not with 50 ng/ml NRG-1 for 10 minutes. Lysates were analyzed by western using beta-tubulin as a loading control. **B)** PC-9 cells expressing ERBB4 variants were treated with the indicated concentrations of osimertinib in the presence or absence of 50 ng/ml NRG-1 for 72 hours and cell viability was measured with the MTT assay. The mean and SD of triplicate analyses of one of 2-3 independent experiments are shown. **C)** PC-9 cells were treated with 100 nM osimertinib for 14 days in the presence or absence of 50 ng/ml NRG-1 and stained with crystal violet. **D)** Quantification of colony area data from C, shown as percentages of DMSO-treated controls. **E)** Colony area fold changes of NRG-1 and osimertinib-treated cells compared to osimertinib alone from the data shown in C and E. P-values were calculated by one-way ANOVA with Šidák’s multiple comparisons test in D and E.

In contrast to the short-term experiments, ERBB4-mutants S303F, E715K and K935I significantly promoted resistance to osimertinib upon prolonged, 14-day treatment (Fig. 6C-D). All three mutants were able to confer osimertinib resistance in the presence of NRG-1, with the E715K-mutant demonstrating the most robust effect. The E715K and K935I-mutants were also able to promote osimertinib resistance in the absence of NRG-1, albeit quite weakly (Fig. 6C-D). These results are consistent with previous observations that activating, cancer-associated ERBB4 mutations typically enhance the ligand-mediated activation of ERBB4 (Fig. 4, (Kurppa *et al*., 2016; Chakroborty *et al*., 2022)). Indeed, both S303F and E715K, as well as the L798R-mutant, demonstrated significantly increased responsiveness to NRG-1 under osimertinib treatment (Fig. 6E).

Together, these results demonstrate that activating ERBB4 mutations are able to confer resistance to EGFR-targeted therapy *in vitro*, and that this ability is particularly evident in the presence of an ERBB4 ligand.

### Activating ERBB4 mutations promote resistance to EGFR-targeted therapy *in vivo*

In order to assess whether the activating ERBB4 mutations can also promote resistance to osimertinib *in vivo*, we subcutaneously implanted the PC-9 cells expressing wild-type ERBB4 or ERBB4-mutants S303F, E715K, L798R and K935I in NMRI nude mice and analyzed the rate of relapse under continuous osimertinib treatment. While none of the PC-9 xenograft tumors expressing wild-type ERBB4 relapsed during the 189-day treatment, 4 out of 13 E715K tumors, 3 out of 12 S303F tumors, 1 out of 12 L798R tumors and 1 out of 13 K935I tumors relapsed during treatment. This translated to worse relapse-free survival under osimertinib for mice bearing ERBB4 E715K or S303F -expressing tumors (E715K, *P* = 0.0398; S303F, *P* = 0.0826) (Fig. 7A-B).

**Figure 7.**
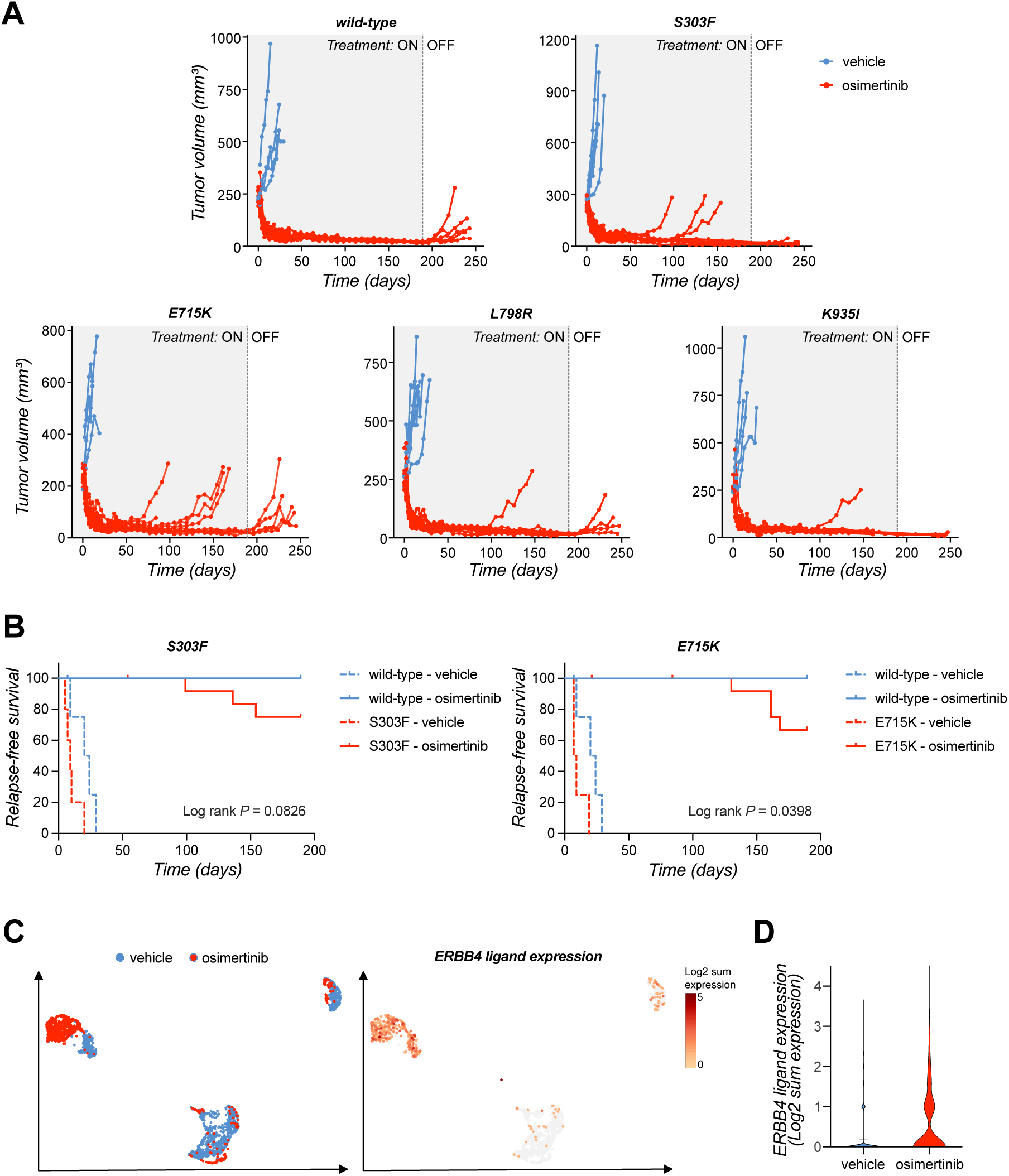
Activating ERBB4 mutations promote resistance to EGFR-targeted therapy *in vivo*. **A)** Tumor volumes of individual mice bearing xenograft tumors of PC-9 cells expressing ERBB4 variants and treated with vehicle (*n* > 4-5 per group) or osimertinib (*n* > 11-13 per group) for 189 days, followed by treatment withdrawal. **B)** On-treatment relapse-free survival of mice treated with vehicle or osimertinib. Log rank test was used for the statistical analysis comparing the survival of mutant or wild-type ERBB4 xenograft-bearing mice under osimertinib treatment. **C)** Single-cell RNA-sequencing analysis of ERBB4 ligand expression in PC-9 xenograft tumors from mice treated with vehicle or osimertinib for 21 days (GSE131604, Kurppa *et al*. 2020). Left: clustering of vehicle-treated (blue) and osimertinib-treated (red) tumor cells in the UMAP space. Right: ERBB4 ligand expression in tumor cells, shown as the log2 sum of *NRG1*, *NRG2*, *NRG3, NRG4*, *HBEGF*, *EREG* and *BTC* expression. **D)** Violin plot of ERBB4 ligand expression in xenograft tumors from vehicle or osimertinib-treated mice.

Considering the observations above that the ability of the ERBB4 S303F and E715K-mutants to confer resistance to osimertinib is at least partly dependent on the availability of ERBB4 ligands (Fig. 6C-E), we analyzed the expression of ERBB4 ligands *in vivo* in PC-9 xenografts using previously published single-cell RNA-sequencing dataset (Kurppa *et al*., 2020). As expected, the vehicle-treated and osimertinib-treated tumor cells clustered mostly separate in the UMAP space, indicating significant transcriptional changes in the tumor cells upon osimertinib treatment (Fig. 7C). The expression of ERBB4 ligands was significantly increased in the osimertinib-treated tumor cells compared to vehicle-treated cells, demonstrating that osimertinib treatment results in upregulation of ERBB4 ligand expression in PC-9 cells *in vivo* (Fig. 7C-D). Thus, it is plausible that this elevation in ERBB4 ligand expression provides the stimulus required for the ERBB4-mutants to promote resistance to osimertinib.

Together, these data provide compelling evidence that activating ERBB4 mutations can promote resistance to EGFR-targeted therapies *in vivo*.

## DISCUSSION

The high overall frequency of somatic *ERBB4* mutations in various cancer types and the existence of clinically approved pan-ERBB inhibitors that potently inhibit ERBB4 have raised the question whether *ERBB4*-mutant cancer patients could benefit from ERBB4-targeted therapy. Understanding the functional consequences of the mutations is indispensable for clinical decision making regarding targeted therapies (Hyman, Taylor and Baselga, 2017). Yet, unlike the frequent *ERBB2* amplifications in breast cancer and the activating kinase domain mutations of *EGFR* in lung cancer, mutations of *ERBB4*, *ERBB2* and *ERBB3* are highly diverse across different cancer types, making it challenging to evaluate their clinical actionability. Especially *ERBB4* has lacked evident hotspot mutations although recurrent *ERBB4* mutations have started to emerge upon increased cancer profiling. While activating, potential driver *ERBB4* mutations have been identified, these are rare in patients, which limits the possibility to clinically evaluate their potential to predict response to ERBB4-targeted therapy (Prickett *et al*., 2009; Kurppa *et al*., 2016; Nakamura *et al*., 2016; Chakroborty *et al*., 2022). In addition, ERBB4 mutations have been detected in patients relapsing on EGFR-targeted therapies (Cremolini *et al*., 2019; Jänne *et al*., 2022; Vokes *et al*., 2022; Yaeger *et al*., 2023), potentially providing another context where ERBB4-targeting might be beneficial. Here, to better facilitate clinical interpretation of cancer-associated *ERBB4* mutations, we functionally characterized 18 recurrent *ERBB4* mutations.

Our results indicated that majority of the analyzed *ERBB4* mutations (11/18) have oncogenic potential. While three of the mutants were strongly transforming in both non-tumorigenic cell models, eight of the analyzed mutants were transforming in just one of the two cell models. Similar cell type-dependent transforming potential has been observed before with other oncogenic ERBB mutations. Strikingly, the most recurrent ERBB3 mutation V104M, known to be oncogenic and frequently occurring in gynecological and gastrointestinal tract cancers, is strongly transforming in MCF10a mammary epithelial cells and colon epithelial cells but not in Ba/F3 lymphoid cells (Jaiswal *et al*., 2013; Koivu *et al*., 2024). Other examples of ERBB4 mutations with cell type-dependent transforming potential include E542K and G741R, as well as R544W that was also analyzed in this study using different cell models than previously (Prickett *et al*., 2009; Chakroborty *et al*., 2022). The underlying causes for such cell type-dependent transforming potential is not fully understood but could be related to the mutant favoring certain dimerization partners or ligands for activating oncogenic signaling, and their different availability in a given cellular context. To overcome the limitations of focusing on a single cellular context, we used two very different cell models to address the transforming potential of ERBB4 variants: MCF10a cells that endogenously express EGFR, ERBB2 and ERBB3, and Ba/F3 cells that endogenously only express low levels of murine ERBB3. It is possible that the remaining 7/18 mutants, not demonstrating significant transforming potential in this study, could be transforming in other cell types or tissue contexts.

Of the three mutants that were strongly transforming in both cell models, E452K has previously been shown to transform NIH-3T3 murine fibroblasts and to promote anchorage-independent growth of SK-MEL-2 human melanoma cells (Prickett *et al*., 2009). In contrast, the only previous functional study of ERBB4 S303F did not detect a growth promoting effect in an *ERBB2*-amplified breast cancer cell line HCC1569 (Elster *et al*., 2018). This could be due to partial redundancy of the growth promoting functions of overactive ERBB2 and ERBB4.

Mechanistic analyses of the three most strongly transforming mutants (S303F, E452K and L798R) showed that they were gain-of-function mutations enhancing the activity of other ERBB receptors and downstream signaling pathways, most prominently S303F. ERBB4 S303F was hyperphosphorylated even in the absence of a ligand, which reflected its ligand-independent transforming potential. Furthermore, our structural analyses proposed that S303F mutation could result in stabilized dimer interactions i) at the dimerization arm interface between the two monomers, and ii) stabilized interactions within a monomer favoring the active conformation. The dimerization assays confirmed that S303F-mutant ERBB4 formed more active homodimers and heterodimers with other ERBB receptors. The mechanism of activation is further supported by similar findings with the paralogous ERBB2 S310F mutation (Diwanji *et al*., 2021). The ERBB2 S310F mutation makes the dimerization arm binding pocket of ERBB2 even more favorable for ERBB3 binding than the respective binding pockets in homodimers of wild-type EGFR or ERBB4, which in turn are considered as the most stable ERBB dimers when bound to their high-affinity ligands (Diwanji *et al*., 2021; Bai *et al*., 2023; Trenker *et al*., 2024). This could perhaps also explain why ERBB4 S303F appears to enhance dimerization and activation of ERBB receptors even in the absence of a ligand.

The higher ERBB3 levels in Ba/F3 and MCF10a cells expressing ERBB4 S303F were indicative of enhanced ERBB3-ERBB4 co-operation, although ERBB4 S303F did not appear to dimerize more with ERBB3 than wild-type ERBB4 in the COS7 cell model. Indeed, knockdown of endogenous ERBB3 indicated that ERBB3 participated in ligand-independent, ERBB4 S303F-mediated transformation of Ba/F3 cells. While the moderate ERBB3 knockdown achieved in the experimental setup precluded definitive conclusions, ERBB3 knockdown did not seem to affect ERBB4-mediated Ba/F3 transformation in the presence of exogenous NRG-1. Moreover, the ERBB3 upregulation is also observed upon Ba/F3 cell transformation with well-established oncogenic mutants of EGFR (L858R, T790M) (Supplementary Fig. S3A) and ERBB2 (S310F) (Koivu *et al*., 2021). This is in line with the recently recognized phenomenon of co-occurring ERBB mutations having synergistic effect on tumor growth and altering therapy sensitivity (Saito *et al*., 2020; Hanker *et al*., 2021). Together, these findings imply that ERBB4 heterodimers with other ERBB receptors can contribute to cell transformation and growth, supporting the rationale for pan-ERBB inhibition approach in targeting mutant ERBB4 in cancer.

Our *in vitro* results together strongly suggest that ERBB4 S303F, E452K and L798R are potential driver mutations, especially S303F that was found to occur usually as a sole ERBB alteration in cancer patient samples. We also showed that the three mutations were targetable with clinically approved pan-ERBB inhibitors neratinib, afatinib and dacomitinib, similar to several previously identified potential driver ERBB4 mutations (Kurppa *et al*., 2016; Nakamura *et al*., 2016; Donoghue *et al*., 2018; Chakroborty *et al*., 2022). Despite no clinical efficacy was observed in the small cohort of seven *ERBB4*-mutant patients treated with neratinib in the SUMMIT trial, all the four gain-of-function mutants found in SUMMIT trial patients were sensitive *in vitro*, and thus, not intrinsically resistant to neratinib. Unfortunately, the small number of *ERBB4* mutant patients does not allow the assessment of the genomic or cancer-type-related factors that were associated with lack of response in *ERBB2* mutant patients. For instance, the SUMMIT trial identified co-occurring *TP53* mutations to associate with a lack of response to neratinib in *ERBB2*-mutant patients (Hyman *et al*., 2018). Interestingly, all the patients with gain-of-function *ERBB4* mutations had a co-occurring *TP53* mutation. Additionally, three out of seven patients with *ERBB4* mutation also had co-occurring driver *ERBB2* mutations, which are known to be sensitive to neratinib in other cancer types but not in the cancer types that these patients had (Chan *et al*., 2016; Hyman *et al*., 2018). As an example, the strong ERBB2 driver mutation S310F is a hotspot in both breast cancer and bladder cancer but only breast cancer patients with S310F mutation responded to neratinib (Hyman *et al*., 2018). We cannot exclude the possibility that such cancer type-specific responsiveness also extends to *ERBB4*-mutant tumors. It is also possible that the *ERBB4* alterations of the SUMMIT trial patients were not strong enough drivers, at least in these cancer types. Considering our strong evidence for ERBB4 S303F being a driver mutation, and the high efficacy of neratinib in breast cancer, in which S303F appears most frequently, its potential as a predictive marker for neratinib should be clinically evaluated.

There is an increasing body of evidence from cell models, xenografts and patient data that ERBB4 can independently and together with ERBB3 (and/or with increased availability of their ligands) compensate for survival and growth signaling upon ERBB2- or EGFR-targeted therapy (Carrión-Salip *et al*., 2012; Wilson *et al*., 2012; Nafi *et al*., 2014; Canfield *et al*., 2015; Yonesaka *et al*., 2015; Donoghue *et al*., 2018; Shi *et al*., 2018; Debets *et al*., 2023). In addition, a recent study showed that *EGFR*-mutant lung cancer patients with co-occurring *ERBB4* mutations have shorter relapse-free survival on osimertinib treatment (Vokes *et al*., 2022), further suggesting that *ERBB4* mutations may be associated with targeted therapy resistance in patients. Our results in the *EGFR*-mutant lung cancer context support this hypothesis, demonstrating that activating ERBB4 mutations are able to promote resistance to EGFR inhibitor osimertinib. Particularly, the S303F and the previously identified, strongly activating E715K mutation were able to confer resistance to osimertinib both *in vitro* and *in vivo*. While wild-type ERBB4 seems to be dispensable for steady-state proliferation for most cancer cell lines, it is intriguing to speculate that mutant ERBB4 activity may provide cancer cells with the means to survive stress associated with a lack of mitogenic signals, such as oncogene-targeted therapy. All the analyzed ERBB4 mutations demonstrated increased activity compared to wild-type ERBB4 upon prolonged serum starvation, and we have previously shown that activating ERBB4-mutants and ligand-stimulated wild-type ERBB4 promotes cell survival upon extended serum deprivation (Sundvall *et al*., 2010; Kurppa *et al*., 2016; Chakroborty *et al*., 2022). These findings point towards the importance of overactive ERBB4 in promoting survival of cancer cells, which aligns well with its role in promoting therapy resistance.

In conclusion, this study demonstrates that recurrent *ERBB4* mutations are potentially actionable in cancer rather than being inconsequential passenger mutations. In addition, we demonstrate that *ERBB4* mutations can promote resistance to targeted therapies. These data provide a rationale for further investigations into treating patients harboring activating *ERBB4* mutations with clinically available second-generation pan-ERBB inhibitors both in the treatment-naïve setting, as well as in the context of acquired targeted therapy resistance.

## ACKNOWLEDGEMENTS

We thank Maria Tuominen and Mika Savisalo for skillful technical assistance, the bioinformatics (Jukka V. Lehtonen), drug discovery and chemical biology and structural biology (FINStruct) infrastructure support from Biocenter Finland and CSC IT Center for Science for computational infrastructure support at the Structural Bioinformatics Laboratory, Åbo Akademi University, the NordForsk Nordic POP (Patient Oriented Products), NordicPharmaTrain projects and the Solutions for Health strategic area of Åbo Akademi University. The animal study was carried out with the support of Turku Center for Disease Modeling (TCDM), University of Turku, Finland; a member of Biocenter Finland.

This study was supported by a research agreement with PUMA technology (K.E., K.J.K), as well as research grants from the Research Council of Finland under grant numbers 346656 (K.J.K), 338466 (K.J.K), 316796 (K.E.), Sigrid Jusélius Foundation (K.J.K, K.E., T.T.A., M.S.J), Jane and Aatos Erkko Foundation (K.J.K), Finnish Cultural Foundation (K.J.K, V.K.O), Cancer Foundation Finland (K.E.), Turku University Hospital (K.E.), InFLAMES Flagship Programme of the Academy of Finland (decision number: 337531) (T.T.A., M.S.J), the European Union – NextGenerationEU instrument and by the Research Council of Finland under grant number 352823 (T.T.A., M.S.J), Cancer Society of South-West Finland (V.K.O), Instrumentarium Science Foundation (V.K.O), K. Albin Johansson Foundation (V.K.O), Orion Research Foundation (V.K.O), and University of Turku Graduate School (V.K.O).

