## Supplementary Data for "Recurrent cancer-associated ERBB4 mutations are transforming and confer resistance to targeted therapies"

### Supplementary Table S1

| Plasmids generated by Gateway cloning | | | | | |
|---|---|---|---|---|---|
| **Plasmid Name** | **Protein change** | **cDNA change** | **Donor vector** | **Destination vector** | **Note** |
| pBABEpuro-gateway-ERBB4 R106C | R106C | c.316C>T | Commercially ordered from Genewiz | pBABEpuro-gateway (Addgene plasmid #51070; Greulich et al. 2012) | |
| pBABEpuro-gateway-ERBB4 S303F | S303F | c.908C>T | | | |
| pBABEpuro-gateway-ERBB4 R393W | R393W | c.1177C>T | | | |
| pBABEpuro-gateway-ERBB4 E452K | E452K | c.1354G>A | | | |
| pBABEpuro-gateway-ERBB4 R524C | R524C | c.1570C>T | | | |
| pBABEpuro-gateway-ERBB4 R544W | R544W | c.1630C>T | | | |
| pBABEpuro-gateway-ERBB4 R711C | R711C | c.2131C>T | | | |
| pBABEpuro-gateway-ERBB4 G741E | G741E | c.2222G>A | | | |
| pBABEpuro-gateway-ERBB4 S774G | S774G | c.2320A>G | | | |
| pBABEpuro-gateway-ERBB4 L798R | L798R | c.2393T>G | | | |
| pBABEpuro-gateway-ERBB4 V840I | V840I | c.2518G>A | | | |
| pBABEpuro-gateway-ERBB4 R847H | R847H | c.2540G>A | | | |
| pBABEpuro-gateway-ERBB4 G870R | G870R | c.2608G>A | | | |
| pBABEpuro-gateway-ERBB4 G907E | G907E | c.2720G>A | | | |
| pBABEpuro-gateway-ERBB4 R992C | R992C | c.2974C>T | | | |
| pBABEpuro-gateway-ERBB4 K1223T | K1223T | c.3668A>C | | | JM-a CYT-2 numbering: K1207T |
| pBABEpuro-gateway-ERBB4 S1289A | S1289A | c.3865T>G | | | JM-a CYT-2 numbering: S1273A |
| pBABEpuro-gateway-ERBB4 R1304W | R1304W | c.3910C>T | | | JM-a CYT-2 numbering: R1288W |

| Plasmids generated by ligation of oligonucleotides | | | | |
|---|---|---|---|---|
| **Plasmid Name** | **Insert** | **Forward oligo** | **Reverse oligo** | **Destination vector** |
| Tet-pLKO-neo-shErbb3 | TRCN0000023432 | CCGGGTTGGATGATTGACGAGAATACTCGAGTATTCTCGTCAATCATCCAACTTTTTG | AATTCAAAAAGTTGGATGATTGACGAGAATACTCGAGTATTCTCGTCAATCATCCAAC | Tet-pLKO-neo (Addgene plasmid #21916; Wiederschain et al. 2009) |

#### Supplementary Figure 1

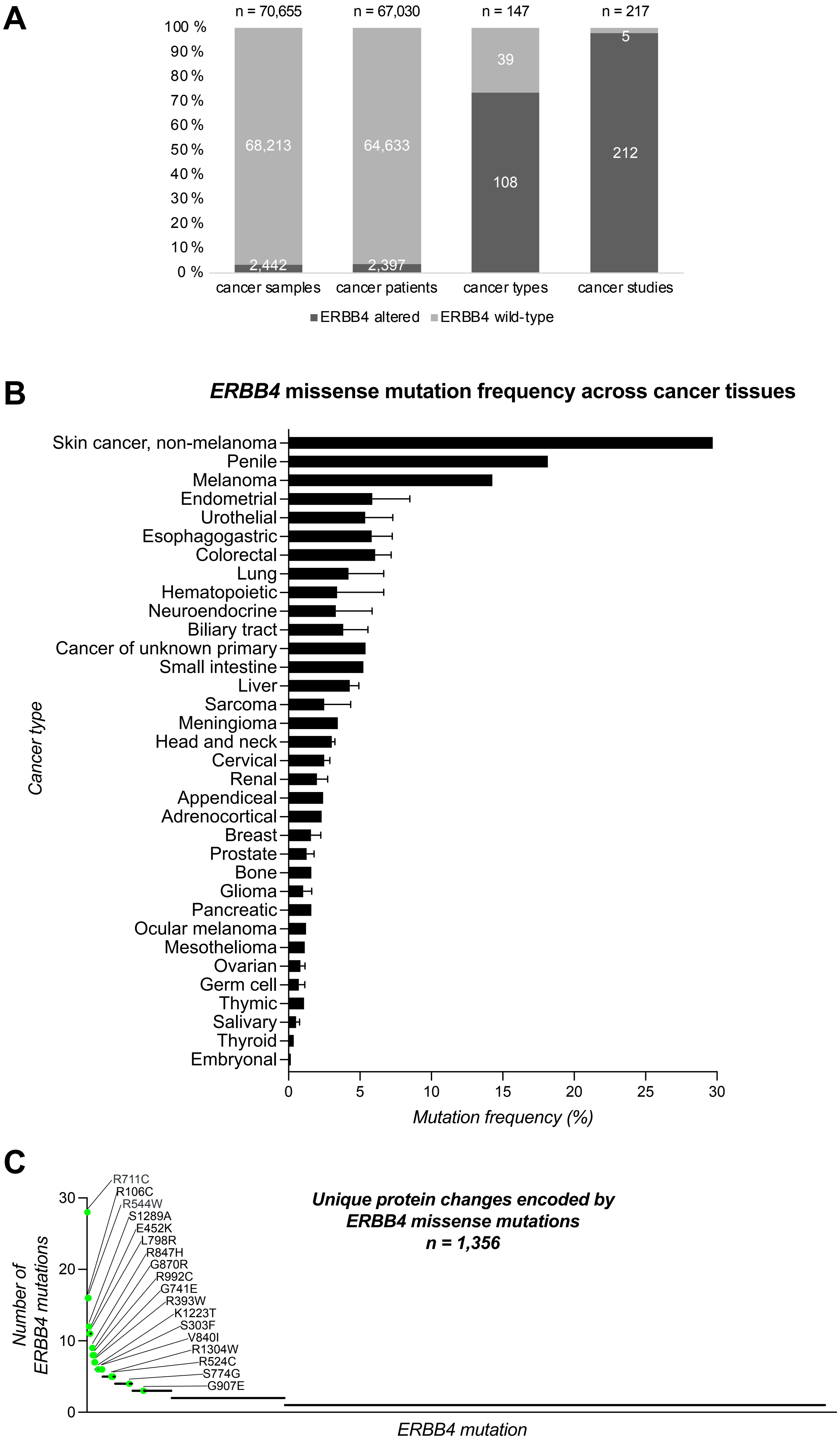

### Supplementary Figure 2

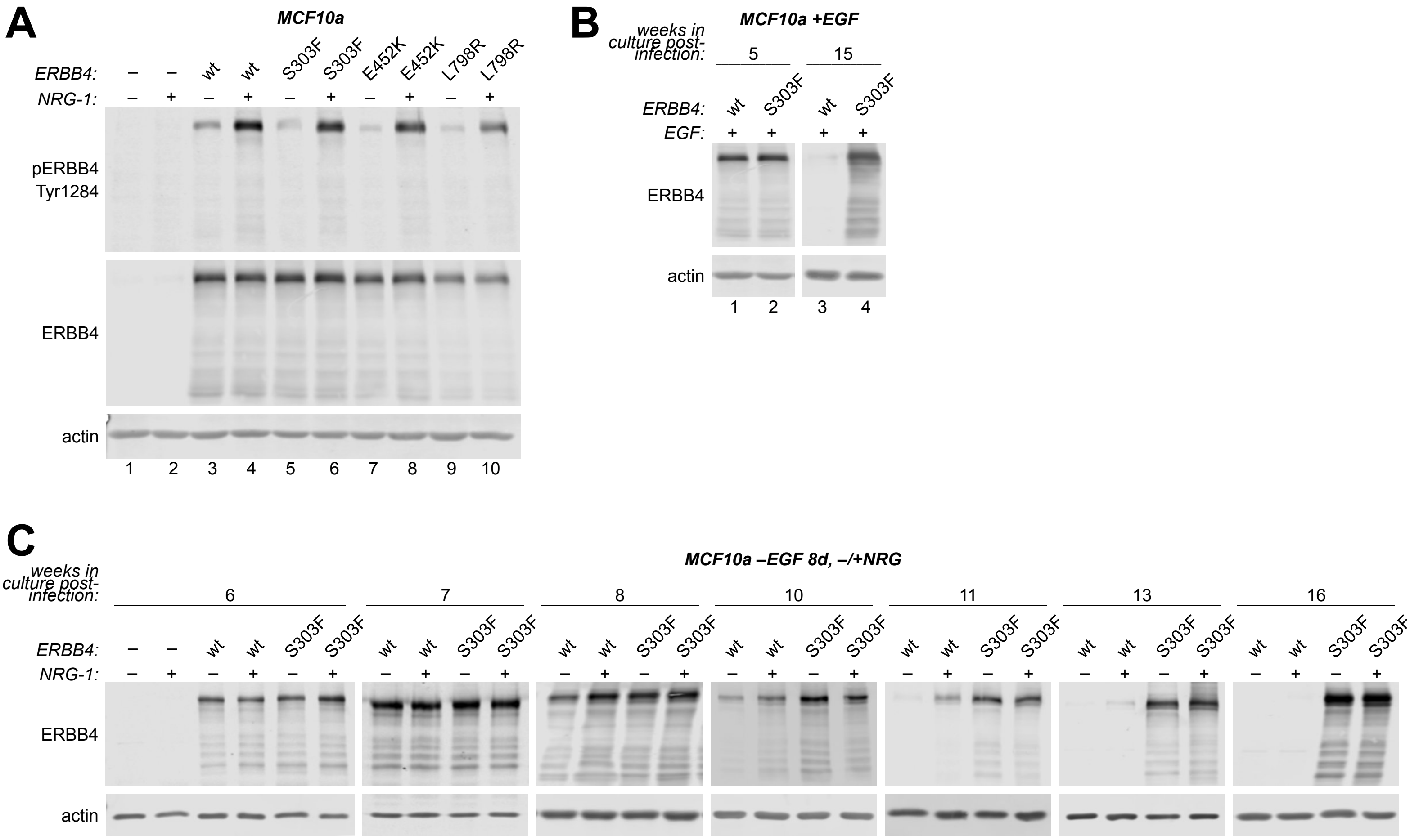

### Supplementary Figure 3

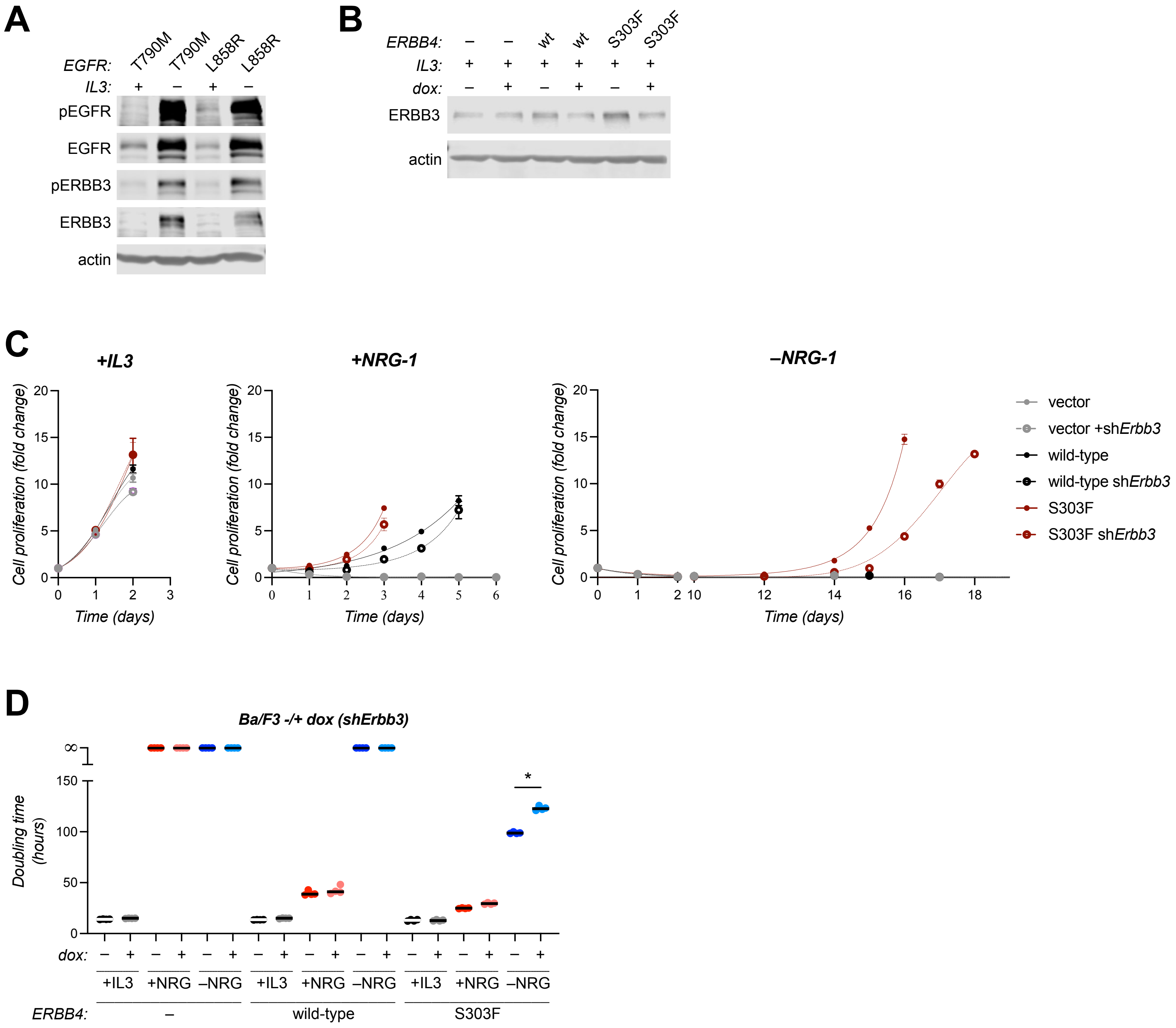

### Supplementary Figure 4

**A**

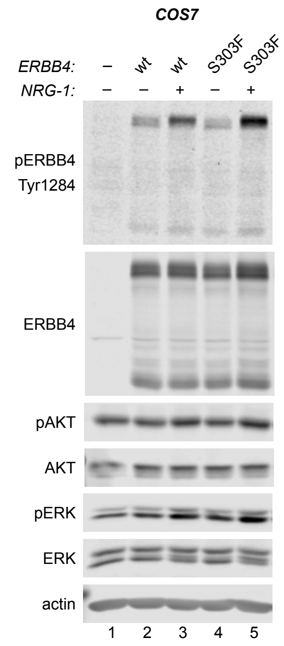

**B**

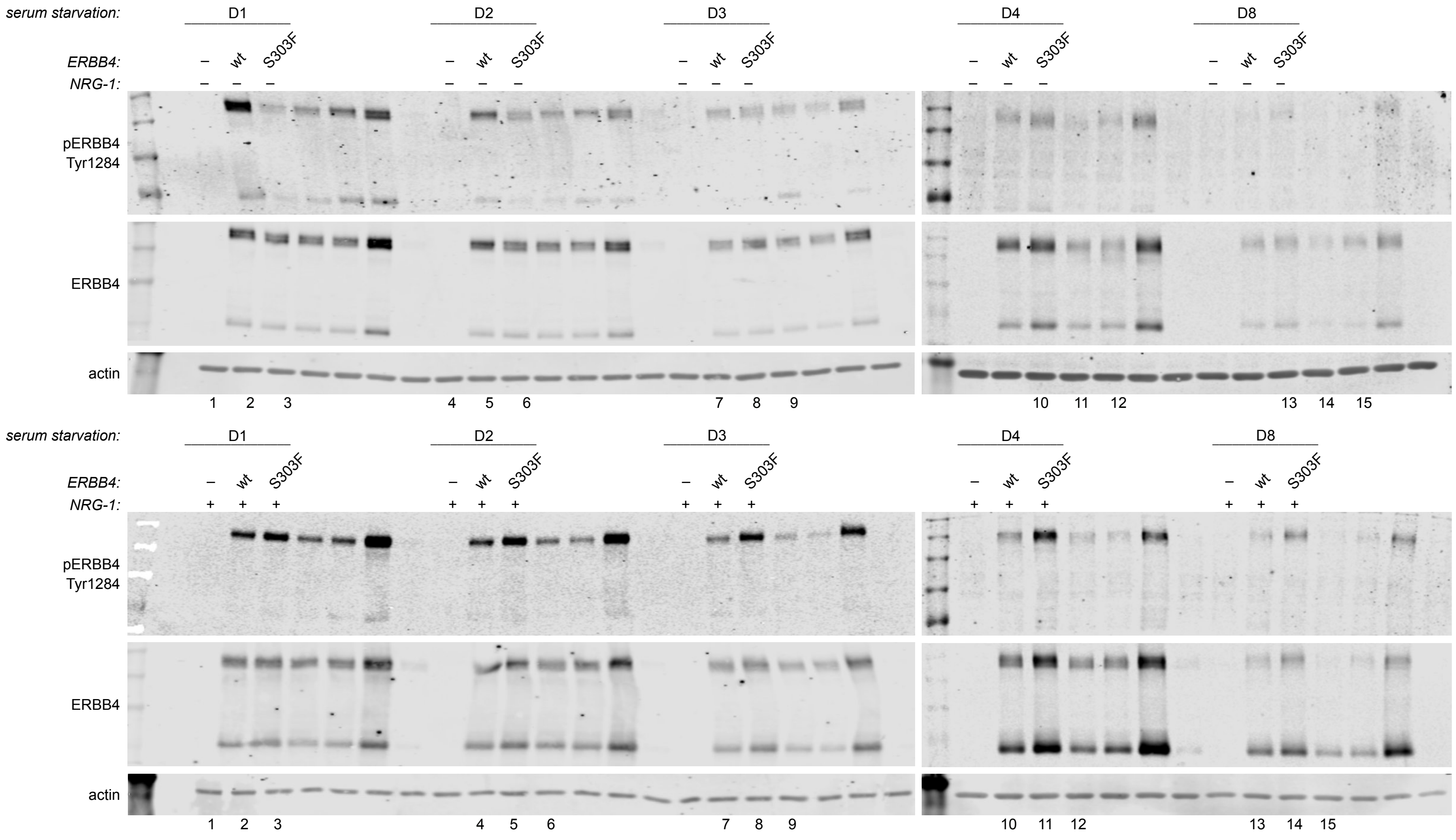

### Supplementary Figure 5

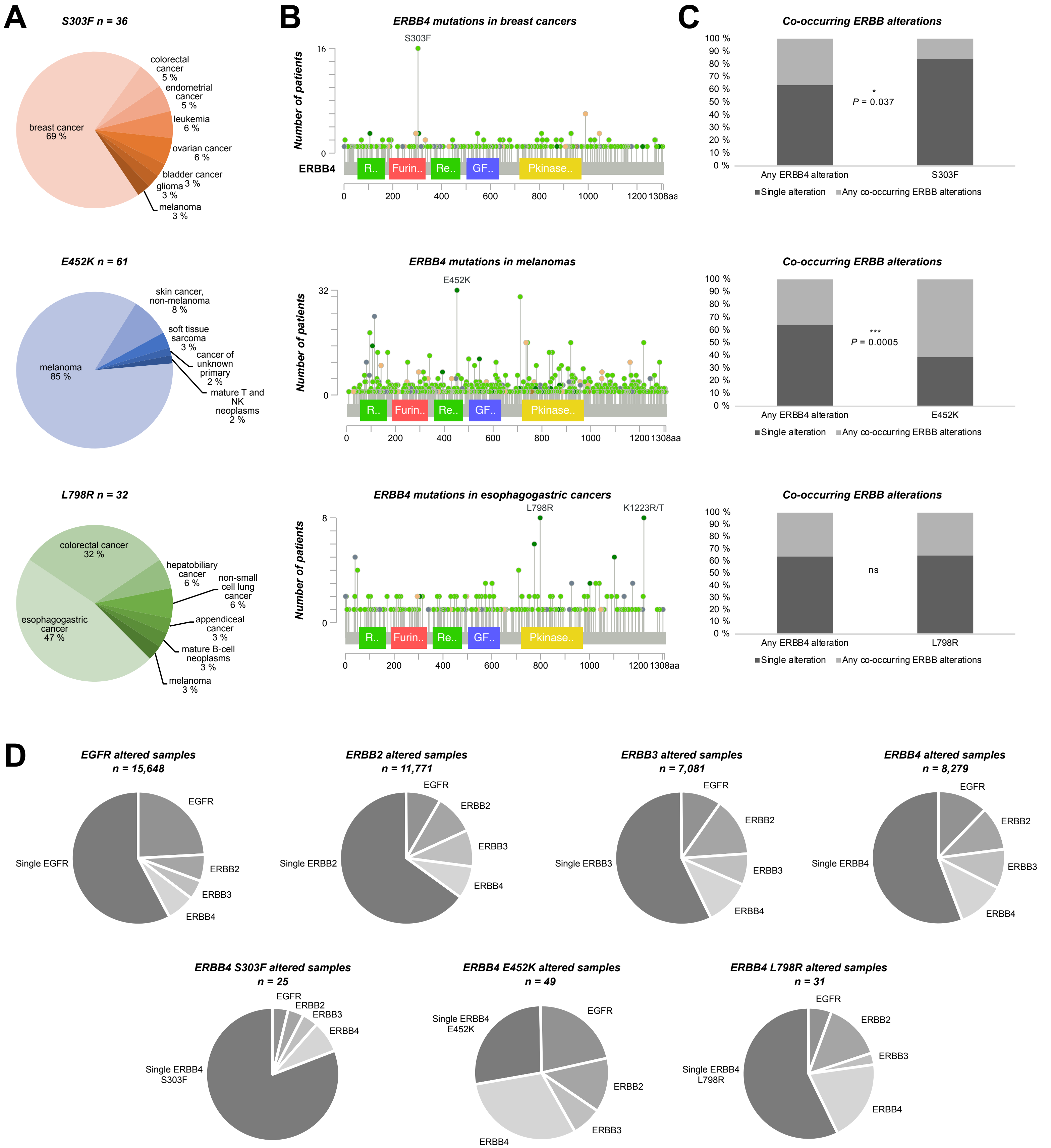

#### SUPPLEMENTARY FIGURE LEGENDS

**Supplementary Figure S1. *ERBB4* alterations in clinical cancer samples. A)** *ERBB4* alterations in clinical cancer samples reported in cBioPortal (https://cbioportal.org) curated non-redundant studies (January 2024). **B)** *ERBB4* missense mutation frequencies across cancer types reported in cBioPortal, classified by tissue of origin. **C)** *ERBB4* missense mutations listed in cBioPortal, ranked by recurrence. The 18 recurrent mutations chosen for functional analyses are indicated.

**Supplementary Figure S2. Biochemical activity and expression of ERBB4 variants in MCF10a cells in the absence of transformation pressure. A)** MCF10a cells stably expressing ERBB4 variants or vector control (-) cells were serum starved overnight and stimulated or not with 50 ng/ml NRG-1 for 10 minutes. Cells were analyzed by western blot using beta-actin as a loading control. **B)** MCF10a cells stably expressing ERBB4 variants or vector control were maintained in fully supplemented growth medium (including 20 ng/ml EGF), and in the presence of puromycin selection for indicated time after lentiviral infection of the cells with the ERBB4 encoding constructs. Cells were analyzed as in A. **C)** MCF10a cells stably expressing ERBB4 variants or vector control were subjected to eight-day EGF deprivation in the presence or absence of 50 ng/ml NRG-1 after an indicated time in culture after lentiviral infection. Cell lysates were analyzed as in A.

**Supplementary Figure S3. Role of ERBB3 in ERBB4-mediated transformation of Ba/F3 cells. A)** Ba/F3 cells stably expressing EGFR mutants were cultured in the absence or presence of IL3 and analyzed by western. **B)** Ba/F3 cells stably expressing ERBB4 variants or vector control (-) together with doxycycline (dox)-inducible *Erbb3* shRNA were cultured in the absence or presence of dox and knockdown efficiency was analyzed by western using beta-actin as a loading control. **B)** Ba/F3 cells analyzed in A were cultured in the presence of IL3 or 20 ng/ml NRG-1 or in the absence of both, with or without dox. Cell proliferation was measured with MTT in triplicate. **C)** Doubling times of ERBB4 variant-expressing Ba/F3 cells upon shRNA-mediated *Erbb3* knockdown. Welch two-sample *t* test was used for pairwise comparisons between -/+ dox samples from each IL3-independently growing cell lines expressing ERBB4 variants. P-values were corrected for multiple comparisons by Bonferroni method. *, $P < 0.0001$.

**Supplementary Figure S4. Effect of prolonged serum starvation on ERBB4 wild-type and S303F mutant activity in COS7 cells. A)** COS7 cells were transfected with constructs encoding ERBB4 variants or vector control (-), then subjected to overnight serum starvation the next day and stimulated or not with 50 ng/ml NRG-1 for 10 minutes. Lysates were analyzed by western blot and loading was controlled with anti-actin. **B)** COS7 cells were transfected with constructs encoding ERBB4 variants or vector control. Next day, the cells were subjected to serum starvation in the presence or absence of 50 ng/ml NRG-1 for 1, 2, 3, 4 or 8 days, and lysates were analyzed as in A.

**Supplementary Figure S5. Tumor characteristics of patients harboring somatic transforming ERBB4 mutations. A)** ERBB4 S303F, E452K and L798R cancer type distribution in cBioPortal (https://cbioportal.org) curated non-redundant studies, AACR GENIE (https://genie.cbioportal.org) and COSMIC (https://cancer.sanger.ac.uk) (redundant data removed). **B)** Lollipop diagram of ERBB4 mutations in the cancer types in which S303F, E452K and L798R are most frequently reported in (diagram sourced from AACR GENIE). **C)** The frequency of co-occurring *ERBB* alterations in patient samples harboring ERBB4 S303F, E452K or L798R mutation, compared to patient samples harboring any ERBB4 alteration. Fisher's exact test was used to compare the ratios. * $P < 0.05$, ** $P < 0.01$, *** $P < 0.001$. Data containing protein coding mutations, copy number alterations and structural variants were sourced from cBioPortal and AACR GENIE (redundant data removed and samples not profiled for all *ERBB* gene alterations excluded). **D)** Co-occurring *ERBB* alterations in cancer samples harboring either *EGFR*, *ERBB2*, *ERBB3*, or *ERBB4* alterations.

#### SUPPLEMENTARY METHODS

##### Lentiviral-mediated *Erbb3* knockdown in Ba/F3 cell lines

Doxycycline-inducible murine *Erbb3* shRNA (shErbb3; TRCN0000023432) expression construct was generated by ligating oligonucleotides listed in Supplementary Table S1 into *Tet-pLKO-neo* vector (a gift from Dmitri Wiederschain: Addgene plasmid #21916; http://n2t.net/addgene:21916; RRID:Addgene_21916) (Wiederschain *et al.*, 2009). The construct was verified by sequencing the insert.

For lentiviral shRNA transduction, HEK293T packaging cells were transfected with lentiviral packaging plasmids (pMLDg/pRRE, pMD2.G, and pRSV-Rev) and *Tet-pLKO-neo-shErbb3* as described in main text and viral supernatants were used to infect Ba/F3 cells. Stable cell pools were selected with 500 μg/ml neomycin (Geneticin, Gibco) for five days and then maintained in 250 μg/ml neomycin. Ba/F3 EGFR L790M and L858R mutant cell lines were generated in our previous study (Chakroborty *et al.*, 2019).

To study the effect of *Erbb3* knockdown on ERBB4-mediated Ba/F3 cell transformation, cells expressing with doxycycline-inducible *Erbb3* shRNA construct were subjected to the transformation assay described in the main text in the presence or absence of 800 ng/ml doxycycline (Millipore). Pairwise comparisons between doubling times of the transformed cells cultured in the presence or absence of doxycycline were made with Welch two-sample $t$ test. P-values were corrected for multiple comparisons by Bonferroni method and $P < 0.0001$ was considered significant.
